# Metabolic rewiring of mitochondria in senescence revealed by time-resolved analysis of the mitochondrial proteome

**DOI:** 10.1101/2022.10.13.512077

**Authors:** Jun Yong Kim, Ilian Atanassov, Frederik Dethloff, Lara Kroczek, Thomas Langer

## Abstract

Mitochondrial dysfunction and cellular senescence are hallmarks of aging. However, the relationship between these two phenomena remains incompletely understood. In this study, we investigated the rewiring of mitochondria upon development of the senescent state in human IMR90 fibroblasts. Determining the bioenergetic activities and abundance of mitochondria, we demonstrate that senescent cells accumulate mitochondria with reduced OXPHOS activity, resulting in an overall increase of mitochondrial activities in senescent cells. Time-resolved proteomic analyses revealed extensive reprogramming of the mitochondrial proteome upon senescence development and allowed the identification of metabolic pathways that are rewired with different kinetics upon establishment of the senescent state. Among the early-responding pathways, the degradation of branched-chain amino acid (BCAA) was increased, while the one carbon-folate metabolism was decreased. Late-responding pathways include lipid metabolism and mitochondrial translation. These signatures were confirmed by metabolic tracing experiments, highlighting metabolic rewiring as a central feature of mitochondria in cellular senescence. Together, our data provide an unprecedentedly comprehensive view on the metabolic status of mitochondria in senescent cells and reveal how the mitochondrial proteome adapts to the induction of senescence.

## Introduction

Cellular senescence (CS) is known to contribute to a wide array of age-related diseases such as cancer, cardiovascular diseases, and osteoarthritis (1). Diverse stressors including genotoxic, epigenotoxic, oxidative, and oncogenic insults induce the senescent state of cells, which is characterized by the secretion of a plethora of bioactive molecules, termed senescence-associated secretory phenotype (SASP) (2). The SASP mainly comprises pro-inflammatory cytokines, growth factors, and extracellular matrix modifiers, which remodel the tissue environment of senescent cells. It largely depends on the type of stress, the type of the recipient cell, and the duration of being senescent. Thus, both the composition and the temporal dynamics of the SASP determine how senescent cells affect their environment. For example, acute SASP is necessary for tissue development and wound healing (3–7), whereas chronic SASP is detrimental and disrupts tissue homeostasis, driving age-related dysfunctions and diseases (8). Accordingly, there has been great interest either in eliminating senescent cells or modulating the chronic SASP to tackle age-related diseases, called senotherapy (9).

Mitochondria have been shown to play regulatory roles in CS and modulate the SASP. Increased mitochondrial biogenesis and decreased turnover of mitochondria by mitophagy results in the accumulation of mitochondria in senescent cells (10). Correlating with the abundance of mitochondria, increased mitochondria-derived reactive oxygen species (mtROS) potentiate the DNA damage response in senescent cells (11) and enhance the SASP by promoting the formation of cytoplasmic chromatin fragments (CCFs), which activate innate immune signaling along the cGAS-STING pathway (12). Moreover, oxidative phosphorylation (OXPHOS) regulates CS and modulates the SASP. Senescent cells are characterized by a higher mitochondrial fatty acid oxidation (FAO) and the inhibition of FAO led to an impaired SASP expression (13). Increased activity of the pyruvate dehydrogenase (PDH) complex, which converts pyruvate to acetyl-CoA in mitochondria, enhances OXPHOS activity and is a rate-limiting factor to drive oncogene-induced senescence (14). Similarly, increased OXPHOS activity in senescent cells governs the strength of the SASP by promoting NAD^+^ regeneration, preventing the activation of AMPK-p53 signaling, which is known to suppress the SASP (15). On the other hand, a decreased cellular NAD^+^/NADH ratio upon OXPHOS dysfunction is sufficient to drive cells into senescence but results in a distinct SASP profile lacking pro-inflammatory IL1 cytokines (16). Together, these studies establish a central role of mitochondria in CS and the SASP and posit mitochondria as an attractive target for senotherapy (17).

Although the importance of mitochondria for CS and the SASP has been established, the functional state of mitochondria in senescent cells remained unclear. Several studies reported an OXPHOS dysfunction and lower mitochondrial membrane potential (MMP) in senescent cells (18–23), whereas the MMP was found to increase with mitochondrial abundance in these cells (24). Moreover, the increased catabolism of central carbons such as pyruvate, fatty acids, and glutamine in senescent cells is difficult to reconcile with dysfunctional mitochondria (13-15, 25-27).

In this study, we performed an in-depth, time-resolved analysis of the mitochondrial proteome upon the establishment of CS. Our findings discover the metabolic rewiring of mitochondria and define their functional status in CS, which provides a possible explanation for apparent discrepancies in the literature.

## Result

### Accumulation of mitochondria with reduced bioenergetic activity in senescent fibroblasts

We treated IMR90 human lung fibroblasts with two chemotherapeutic agents, decitabine and doxorubicin to establish CS. Decitabine is a deoxycytidine analog harboring nitrogen instead of a carbon atom at the 5’ position of the pyrimidine ring (Supplementary figure 1A). Upon incorporation into the replicating genomic DNA, it impairs DNA methylation and causes epigenetic stress. Doxorubicin, on the other hand, blocks topoisomerase II and causes DNA damage. Treatment of IMR90 fibroblasts with decitabine or doxorubicin for 7 days increased mRNA levels of CDKN1A, decreased mRNA levels of LMNB1, and induced the common SASP genes IL1A and IL6, indicating the senescent state (Supplementary figure 1B, 1C). Consistently, the cells had little or no cell-cycle activity (Supplementary figure 1D) and 60-70% of them were senescence-associated β-galactosidase (SA-β-gal) positive (Supplementary figure 1E, 1F). These data demonstrate the successful establishment of the senescent state.

**Figure 1.**
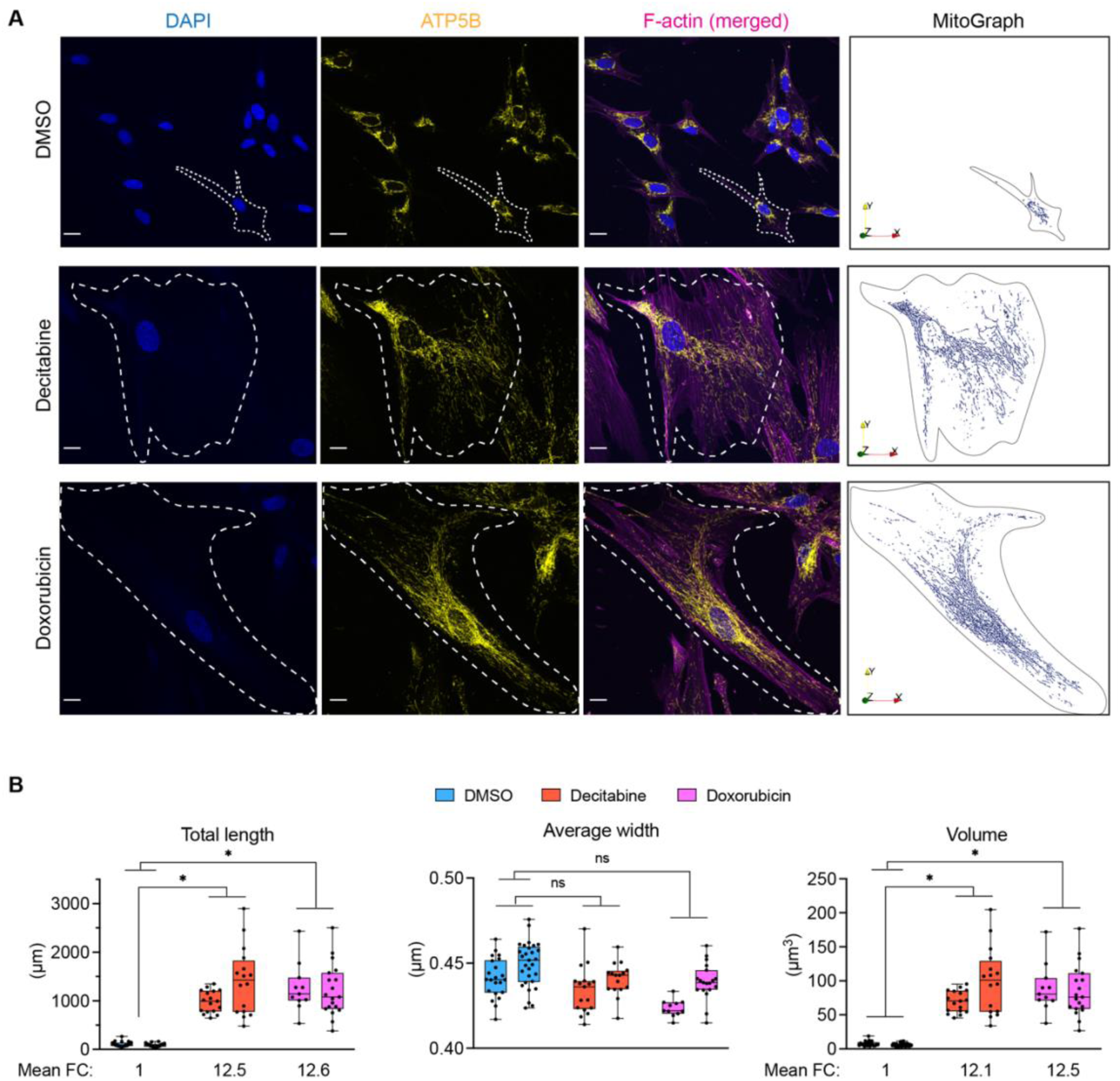
Determination of the mitochondrial volume in senescent fibroblasts. (A) Representative images of mitochondria in IMR90 fibroblasts on day 7 after the treatment with DMSO, decitabine, or doxorubicin. (Left) The maximal projection of confocal images from different z-levels is shown. (Right) Each z-stack image was combined and rendered into a three-dimensional image using MitoGraph 3.0 (29). ATP5B was used as a mitochondrial marker. F-actin was stained to define a cellular boundary. AU: arbitrary unit. (B) Quantification of mitochondrial length, width, and volume using MitoGraph 3.0. Each dot represents a single cell value. Between 11 and 29 cells were analyzed per replicate and condition. Whisker stands for the mean. Mean fold changes are shown. Nested one-way ANOVA, Dunnett correction. n=2.

To examine mitochondrial functions in senescent IMR90 fibroblasts, we first aimed to unambiguously determine the mitochondrial abundance in these cells. We therefore determined the volume of mitochondria, rather than relying on a two-dimensional analysis of the mitochondrial network. We used an immunocytochemistry-based quantification method, which, in contrast to other probes such as MitoTracker and NAO, allowed the determination of the mitochondrial volume largely independent of mitochondrial activities (28). Confocal images of mitochondria from a single cell were stacked and rendered into a three-dimensional image using MitoGraph 3.0 (Figure 1A) (29). It showed that the mitochondrial length was increased on average >12-fold in a senescent fibroblast, while the average width remained unaltered (Figure 1B). Accordingly, the volume of mitochondria was increased over 12-fold in a senescent fibroblast when compared to a proliferating fibroblast. This is largely commensurate with the 8-fold increase in the volume of a senescent fibroblast (30).

We next determined mtDNA levels in senescent IMR90 fibroblasts. Although the relative amount of mitochondrial DNA (mtDNA) was higher in these cells, normalization to the mitochondrial volume revealed decreased mtDNA levels per mitochondrion (Figure 2A). Similarly, the measurement of mitochondrial membrane potential (MMP) with TMRM showed an increase in the MMP in the senescent fibroblasts (Figure 2B-E). However, normalization to the mitochondrial volume revealed a decreased MMP per mitochondrion (Figure 2B-E). In agreement with the alterations in the MMP, senescent cells showed an increased oxygen consumption rate (OCR) on a cellular basis, which however corresponds to a decreased OCR per mitochondrial volume (Figure E, F). We observed a similar pattern for mitochondrial superoxide levels in the senescent fibroblasts (Figure 2G, 2H).

**Figure 2.**
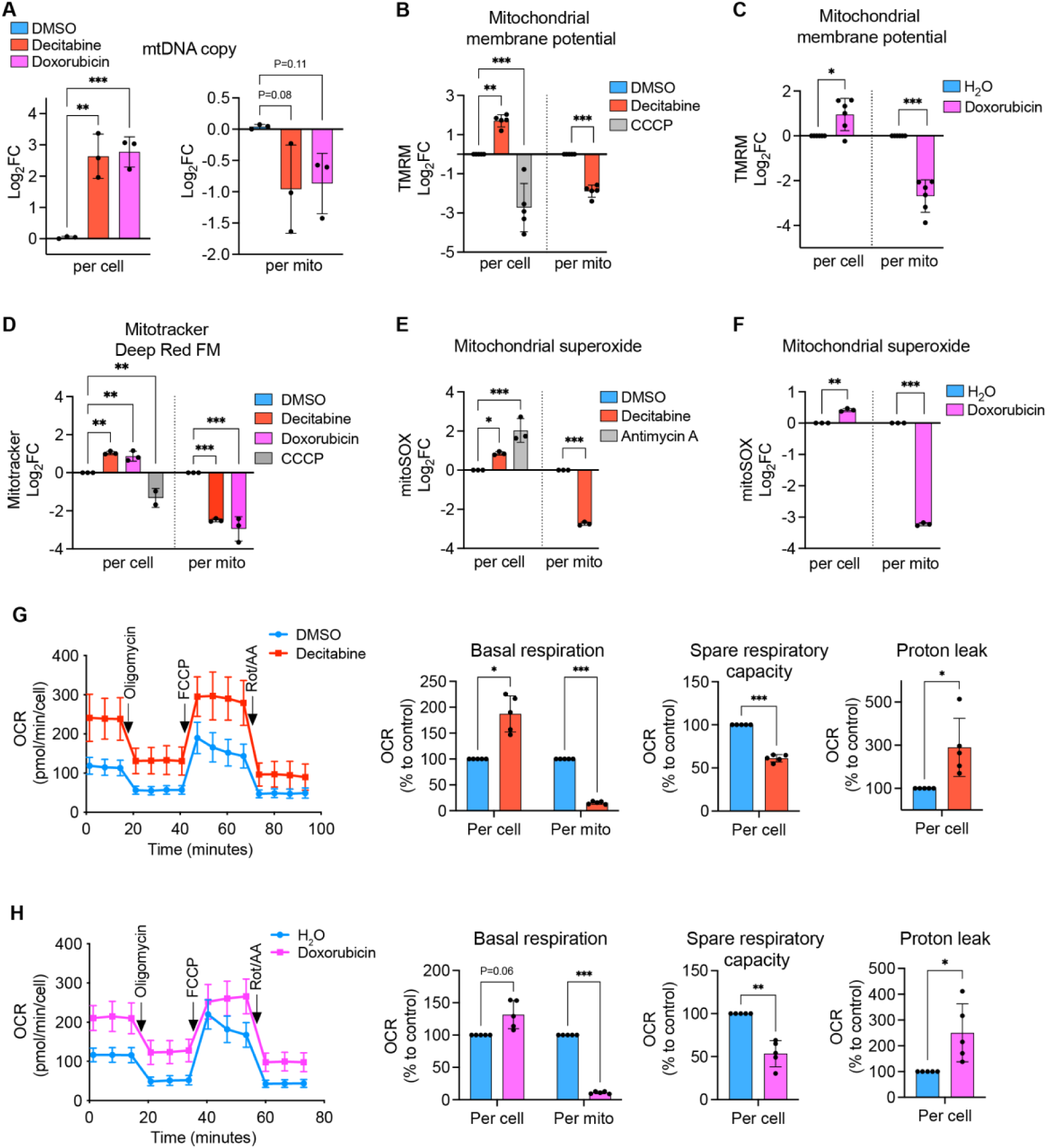
Accumulation of mitochondria with reduced bioenergetic activity in senescent fibroblasts. The values are first measured per cellular basis on day 7 after the treatment with DMSO, decitabine, or doxorubicin to IMR90 fibroblasts and followed by scaling with the relative mitochondrial volume per cell. (A) Determination of mtDNA levels in cellular DNA extracts by RT-qPCR using MT-ND1 and ACTB as probes for mtDNA and nuclear DNA, respectively. One-way ANOVA, Dunnett correction. n=3 from independent cultures. (B-F) Measurement of mitochondrial membrane potential and mitochondrial superoxide level. IMR90 fibroblasts were subjected to staining with TMRM (B, C), Mitotracker Deep Red FM (D), or mitoSOX (E, F) and analyzed by flow cytometry. Antimycin A (10 µM) and CCCP (50 µM) were used as controls. (B, D, E): one-way ANOVA, Dunnett correction. (C, F): Welch t-test, Bonferroni-Dunn correction. (B): n=5, (C): n=6, (D, E, F): n=3 from independent cultures. (G, H) Measurement of oxygen consumption rate (OCR). Left: real-time OCR before and after the sequential addition of oligomycin (1 µM), FCCP (0.5 µM), and rotenone/antimycin A (Rot/AA, 0.5 µM each). Right: Fold changes calculated from the left graphs. The % values were transformed to Log2 values and subjected to statistical analysis. Welch t-test, Bonferroni-Dunn correction. n=5.

We therefore conclude that the bioenergetic activity of mitochondria is decreased in senescent fibroblasts. However, the accumulation of such hypoactive mitochondria results in the overall enhancement of mitochondrial functional parameters in these cells. These findings highlight the importance to consider mitochondrial abundance when assessing the functional status of mitochondria and possibly resolve conflicting reports on mitochondrial fitness in senescent cells.

### Reshaping of the mitochondrial proteome during CS development

To gain further insights into the reprogramming of mitochondria in senescence, we monitored the rewiring of the mitochondrial proteome upon transition to the senescent state by tandem mass tag labeling mass spectrometry (TMT-MS) in a time-resolved fashion (Figure 3A). Cellular proteomes were analyzed on days 1, 3, 5, and 7 after the treatment with DMSO or decitabine (Figure 3A). 6482 proteins were quantified in all samples, corresponding to more than 80% of the total identified proteins, which showcased the power of TMT-MS to minimize missing values across the samples, and hence were used for further analyses (Supplementary figure 2A). Principal component analysis (PCA) revealed progressive changes in the cellular proteome during the development of the CS, while that of proliferating control cells with DMSO remained largely unaltered as expected (Supplementary figure 2B). Changes in protein levels of key senescence markers confirmed the establishment of the CS by decitabine (Supplementary figure 2C). Our dataset covered around 60% of the proteomes of major cellular organelles based on the reference proteome of each organelle, including nucleus (31), cytosol (32), ER membrane (33), and mitochondria (34) (Supplementary figure 2D).

**Figure 3.**
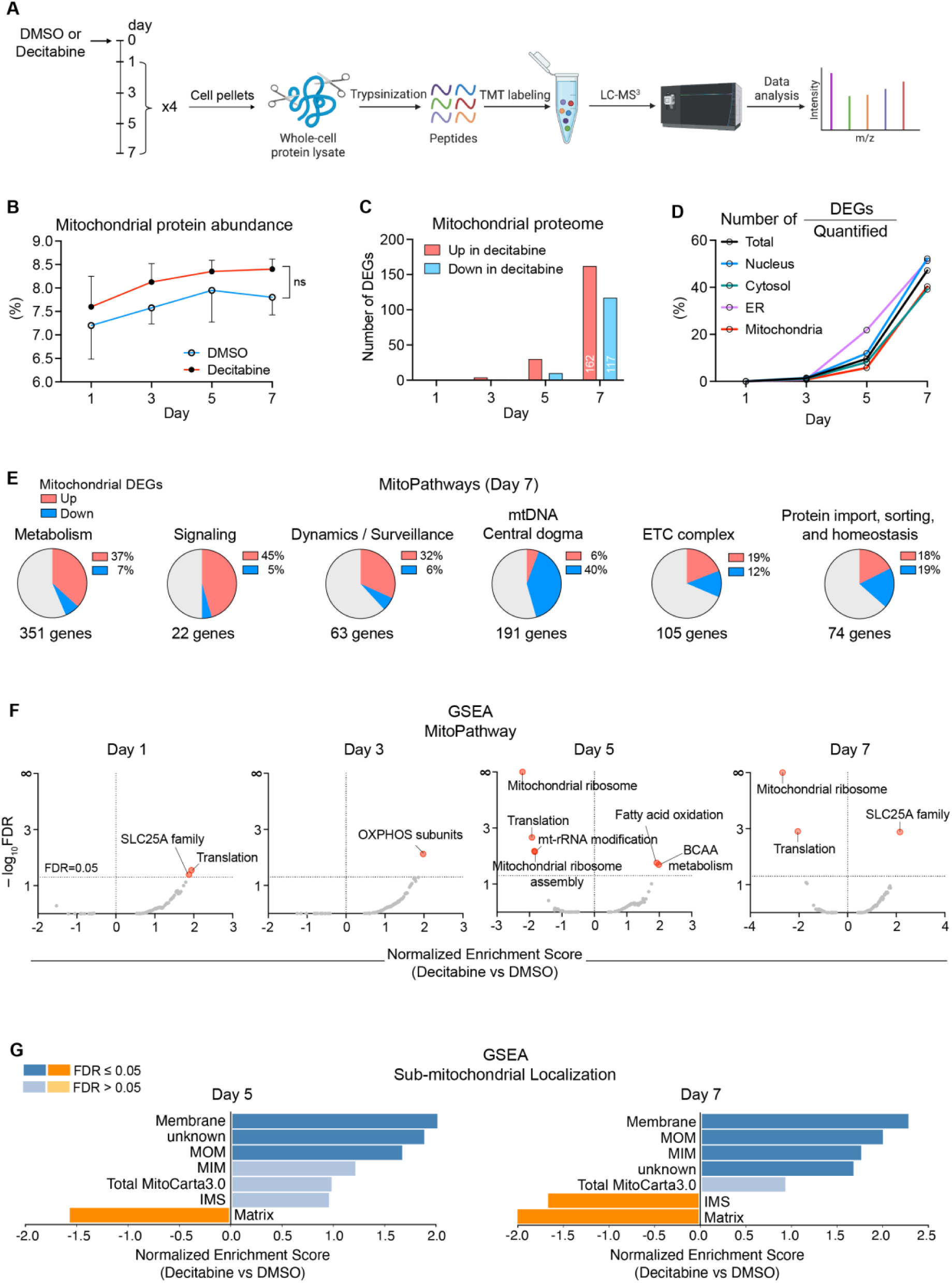
Reprogramming of the mitochondrial proteome upon the development of CS. (A) Workflow for the time-resolved analysis of the mitochondrial proteome upon CS induction by decitabine. Cellular proteome was measured by tandem mass tag labeling mass-spectrometry (TMT-MS) on days 1, 3, 5, and 7 after DMSO or decitabine treatment in IMR90 fibroblasts. All samples were measured in biological quadruplicates. (B) The percentage of mitochondrial proteins within the cellular proteome during CS development. Mitochondrial protein abundance was calculated as 2^Σ(mitochondrial peptides reporter intensities)^/2^Σ(total reporter intensities)^. Welch t-test at each time point, Bonferroni-Dunn correction. n=4. (C) The number of differentially expressed genes (DEGs) encoding mitochondrial proteins at each time point of CS development. (D) The percentage of DEGs encoding different organellar proteins among all quantified proteins at different time points of CS development. (E) Representation of mitochondrial pathways within DEGs on day 7. Six major categories can be distinguished using MitoPathways enlisted in the human MitoCarta 3.0. The number of genes beneath the circles indicates the number of quantified proteins in each category. (F) Gene set enrichment analysis (GSEA) of the proteomics data according to MitoPathways. FDR=0.05 was used as the cutoff. (G) GSEA of sub-mitochondrial localization of the proteomics data on days 5 and 7 according to the human MitoCarta 3.0. MOM: mitochondrial outer membrane, MIM: mitochondrial inner membrane, IMS: intermembrane space.

To define mitochondrial proteomic changes, we first examined whether the increased mitochondrial abundance in senescent cells introduces bias in our proteomic analysis. We calculated the proportion of mitochondrial proteins in the cellular proteome at each time point but did not observe alterations in the fraction of mitochondrial proteins during the CS development, unlike that of ER membrane or nuclear proteins (Figure 3B, Supplementary figure 2F). We also compared mitochondrial proteomic changes by two different normalization units: total peptide counts and mitochondrial-specific peptide counts. The comparison yielded extremely high correlations between fold changes calculated by the two normalization units at all time points (Supplementary figure 2E). These results indicate that the mitochondrial proteome increased proportionately to the cellular proteome throughout the development of the CS and the proteomic changes can be faithfully analyzed from the data normalized by the total peptide counts.

The mitochondrial proteome was significantly altered during the CS development, yielding 279 differentially expressed genes (DEGs) on day 7, corresponding to nearly 40% of the total mitochondrial proteins quantified (Figure 3C, 3D). We observed similar changes for nuclear, cytosolic, and ER membrane proteins, which indicates a lack of strong bias in organellar proteomic changes during the CS development (Figure 3D). Based on the affected mitochondrial pathways (MitoPathways, curated in MitoCarta 3.0), the 279 mitochondrial DEGs on day 7 were categorized into 6 major groups and the percentage of DEGs within each group was calculated. This analysis showed a general upregulation of genes related to metabolism, signaling, dynamics/surveillance, and downregulation of mtDNA-related genes, whereas we observed mixed alterations in genes related to OXPHOS and mitochondrial proteostasis (Figure 3E). Changes in the mitochondrial proteome were also subjected to a gene set enrichment analysis (GSEA), which highlighted alterations in metabolic pathways (e.g. branched-chain amino acid metabolism, fatty acid oxidation, SLC25A family) and in the translation of mtDNA-encoded genes (Figure 3F), corroborating the previous analysis (Figure 3E). Moreover, the GSEA of sub-mitochondrial localization revealed a general increase in inner and outer membrane proteins, while matrix proteins were decreased on day 5 and 7 (Figure 3G). These alterations mainly result from a general increase in SLC25A family proteins which are integral membrane proteins and an overall decrease in the translational apparatus in the matrix space. The increase in membrane proteins is unlikely due to the enhanced protein import because the small TIM proteins (TIMM8B, TIMM9, TIMM10, TIMM13) which are responsible for the chaperone-mediated import of many hydrophobic membrane proteins were reduced altogether on day 7 (Supplementary figure 3; protein import & sorting). Considering that the mitochondrial protein abundance remained constant throughout the CS development, these analyses suggest remodeling of the mitochondrial proteome with an altered ratio between the membrane and matrix proteins.

### Changes in the mitochondrial proteome reveal broad metabolic rewiring of senescent fibroblasts

We further monitored how individual pathways were affected throughout the CS establishment. We identified four groups of DEGs differing in their temporal dynamic patterns (Figure 4A, Supplementary figure 4 for detailed lists of genes). Groups 1 and 2 include mitochondrial proteins that were altered rather early (day 3 or 5) and whose abundance either kept increasing (group 1) or decreasing (group 2) upon the decitabine treatment (Figure 4A). On the contrary, groups 3 and 4 contain late-responding proteins, whose abundance changed only on day 7. Notably, only 1.5 % of the analyzed DEGs fluctuated over the CS development and did not fall into any of the four groups (Figure 4C). Each group of proteins was then subjected to an over-representation analysis with KEGG and GO databases (Figure 4B). The analysis revealed an enrichment of branched-chain amino acid (BCAA) catabolism in group 1 as well as the mitochondrial arm of the one-carbon (1C)-folate metabolism in group 2 (Figure 4B). Proteins associated with fatty acid metabolism and calcium import into mitochondria were over-represented in the late-responding group 3, indicating upregulation of these pathways upon senescence induction (Figure 4A, B), in agreement with previous findings (13, 35). On the contrary, we observed a strong enrichment of subunits of mitochondrial ribosomes and components of the mitochondrial gene expression apparatus in group 4, suggesting a reduction of mitochondrial translation in the established senescent state (Figure 4A, B). To validate these findings and to exclude any bias due to mitochondrial abundance, we synthesized mtDNA-encoded proteins in isolated mitochondria in the presence of ^35^S-methionine (Figure 4D). In agreement with our proteomic analysis, mitochondrial translation was reduced in the senescent fibroblasts (Figure 4D, 4E).

**Figure 4.**
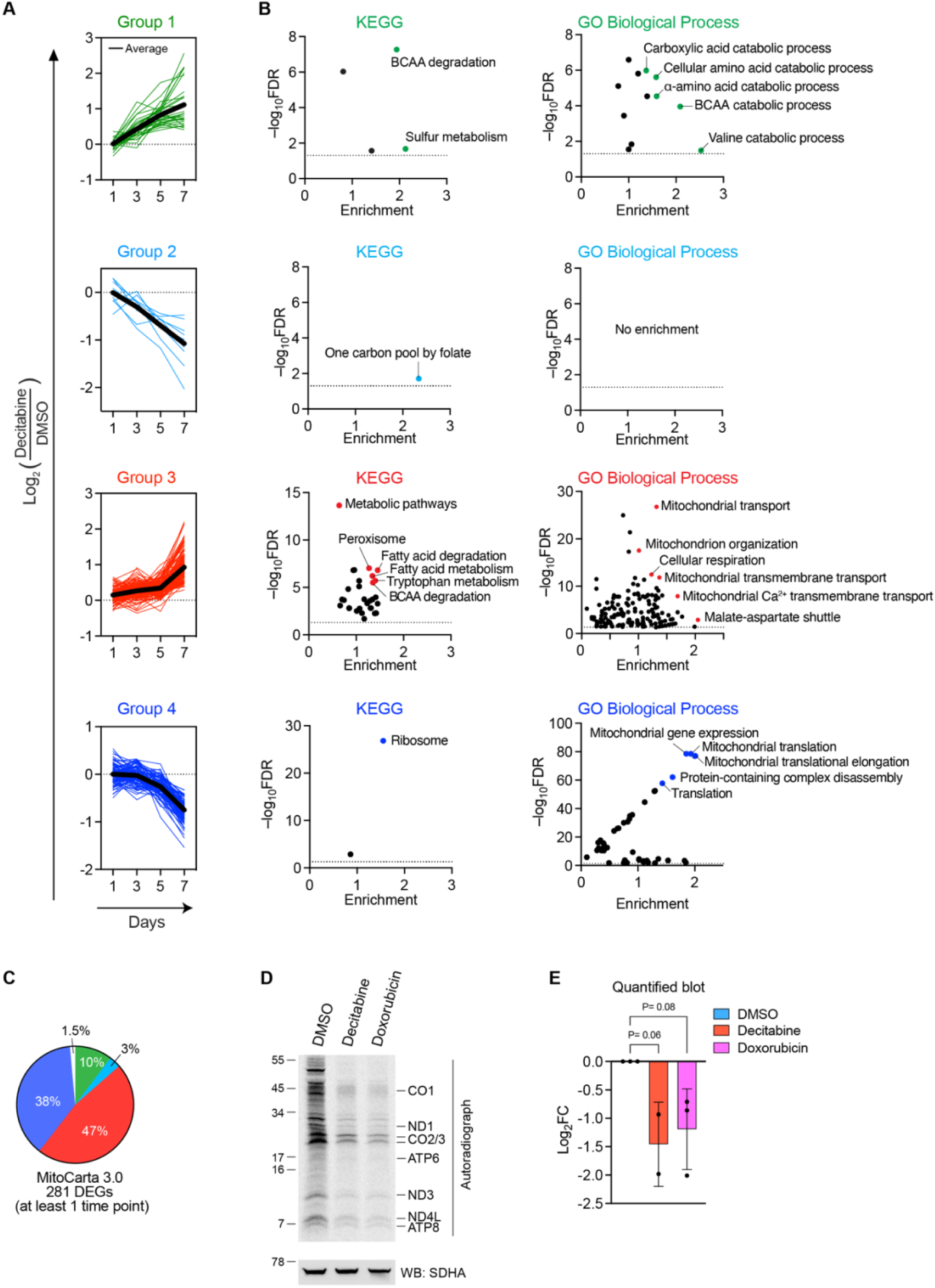
Classification of mitochondrial proteins according to their time-dependent changes during the CS development. (A) Classification of mitochondrial DEGs into four groups according to the temporal dynamics of Log2 fold changes during the development of CS. The bold line in each group indicates the average of Log2 values. For detailed lists of genes in each category, see Supplementary figure 4. (B) Over-representation analysis of mitochondrial DEGs in each group from (A) based on two different databases. KEGG: Kyoto Encyclopedia of Genes and Genomes, GO: Gene Ontology. Highlighted are terms with the highest enrichments and/or significance. The color corresponds to the groups in (A). (C) Percentage of each group within total mitochondrial DEGs from (A). The color corresponds to the groups in (A). (D) *In organello* assay of mitochondrial translation. Mitochondria were isolated from IMR90 fibroblasts on day 7 after the treatment with indicated compounds and incubated in a translation buffer in the presence of ^35^S-methionine as described in Material and Methods. Each protein was annotated based on direct verification by immunoblot or size information. SDHA blot was used as a reference for equal loading. (E) Quantification of (D). Radioactivity in each total lane was quantified and divided by the intensity of the SDHA blot. Log2 fold change relative to the DMSO-treated control is shown. One-way ANOVA, Dunnett correction. n=3 for decitabine, n=2 for doxorubicin.

Together, we conclude that mitochondria are broadly rewired upon CS induction, pointing to metabolic adaptations, favoring the degradation of BCAA but downregulating the 1C-folate metabolism. This is accompanied by a decrease in mitochondrial translation, consistent with the observed decreased respiratory activity of mitochondria in senescent cells.

### Enhanced catabolism of BCAA in senescent fibroblasts

In further experiments, we used metabolic tracing experiments to validate early metabolic adaptations indicated by our proteomic analysis. We observed the accumulation of enzymes of the BCAA metabolism upon CS induction, suggesting an enhanced degradation of BCAAs in the senescent cells (Figure 5A). The nitrogen in BCAAs accumulates in glutamate, which is used to synthesize several non-essential amino acids (NEAAs). On the other hand, carbon atoms of BCAA are found in acyl-CoAs, used for the synthesis of fatty acids or cholesterol, or fed into the TCA cycle (Figure 5B). To monitor the catabolism of BCAA in senescent cells, we performed metabolic tracing experiments with BCAAs that are labeled with stable isotopes of either nitrogen or carbons. These experiments revealed an increased flux of both carbons and nitrogen to the downstream metabolites in the senescent fibroblasts (Figure 5C, 5D). We also observed an accumulation of BCAAs-derived short-chain acylcarnitines such as acetyl-carnitine, propionyl-carnitine, and isobutyryl-carnitine (Supplementary figure 5C), which in agreement with the observed respiratory deficiency and points to enhanced BCAA degradation (36). These data demonstrate the validity of the proteomics signature of increased BCAA degradation in senescent cells.

**Figure 5.**
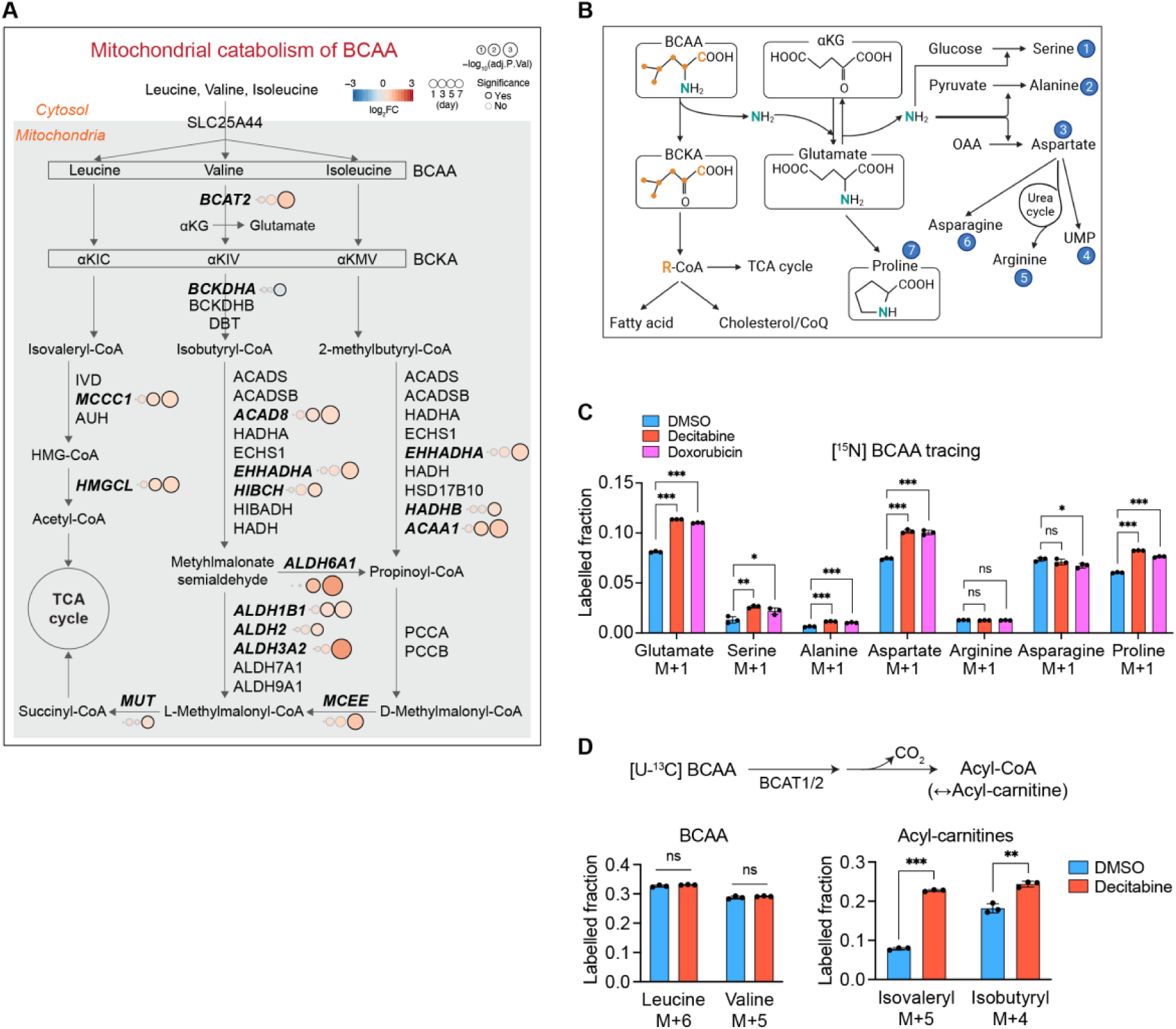
Enhanced mitochondrial BCAA degradation in senescent fibroblasts. (A) The BCAA catabolism in mitochondria. Enzyme levels are derived from the proteomics data. The DEGs are shown as closed circles and in bold italicized font. The color code indicates Log2 fold changes. BCKA: branched-chain α-keto acid, αKIC: α-keto-isocaproate, αKIV: α-keto-isovalerate, αKMV: α-keto-beta-methylvalerate. (B) Carbon and nitrogen flux throughout BCAA catabolism. Metabolites incorporating BCAA-derived nitrogen are marked with circled numbers and highlighted in blue. αKG: α-ketoglutarate, CoQ: coenzyme Q, OAA: oxaloacetate. (C) Metabolic tracing of the BCAA metabolism using ^15^N-L-leucine and ^15^N-L-valine at equimolar concentrations (100 µM each) for 24 h in IMR90 fibroblasts on day 7 after the treatment of indicated compounds. One-way ANOVA for each metabolite, Dunnett correction. n=3 from independent cultures. (D) Metabolic tracing of the BCAA metabolism using ^13^C_6_-L-leucine and ^13^C_5_-L-valine at equimolar concentrations (100 µM each) for 2.5 h in IMR90 fibroblasts on day 7 after the treatment of indicated compounds. Welch t-test, Bonferroni-Dunn correction. n=3 from independent cultures.

### Early reduction of 1C-folate metabolism in senescent fibroblasts

Our proteomic analysis also suggested that the 1C-folate cycle is an early-responding pathway that is rapidly reduced upon the decitabine treatment (Figure 4B; group 2, Supplementary figure 4A; group 2). Notably, although the over-representation analysis was restricted to mitochondrial proteins, enzymes involved in the cytosolic arm of 1C-folate metabolism were also acutely decreased upon induction of the senescent state (Figure 6A). To validate the proteomic footprints in 1C-folate metabolism, we performed targeted metabolomics, focusing on polar metabolites including nucleotides and amino acids. PCA showed that the metabolome of the senescent fibroblasts is distinct from that of proliferating cells (Supplementary figure 5A). The metabolomics revealed a significant reduction of purines (AMP, GMP) and deoxythymidines (dTTP, note that dTMP was under the detection threshold exclusively in the senescent cells), which is indicative of the reduced 1C-folate metabolism (Figure 6B, Supplementary figure 5B, 5C). Another indicator of the activity of the pathway is the serine catabolism by cytosolic SHMT1 and mitochondrial SHMT2. We found decreased glycine levels and an increased serine-to-glycine ratio in the senescent cells, consistent with the decreased SHMT2 level in these cells (Figure 6C, D). These observations were further substantiated by tracing carbons of glucose, which demonstrated that the formation of serine from glucose and glycine from serine was significantly reduced (Figure 6E). To distinguish effects on the cytosolic and mitochondrial arm of the 1C-folate metabolism, we performed tracing experiments using a serine stable isotope with deuterium (Figure 6F). Monitoring the accumulation of dTTP isotopologues allowed us to determine the directionality of the pathway (37). M+1 dTTP was exclusively detected but not M+2 dTTP (Figure 6G), indicating that serine was catabolized exclusively in the mitochondria in proliferating IMR90 fibroblasts.

**Figure 6.**
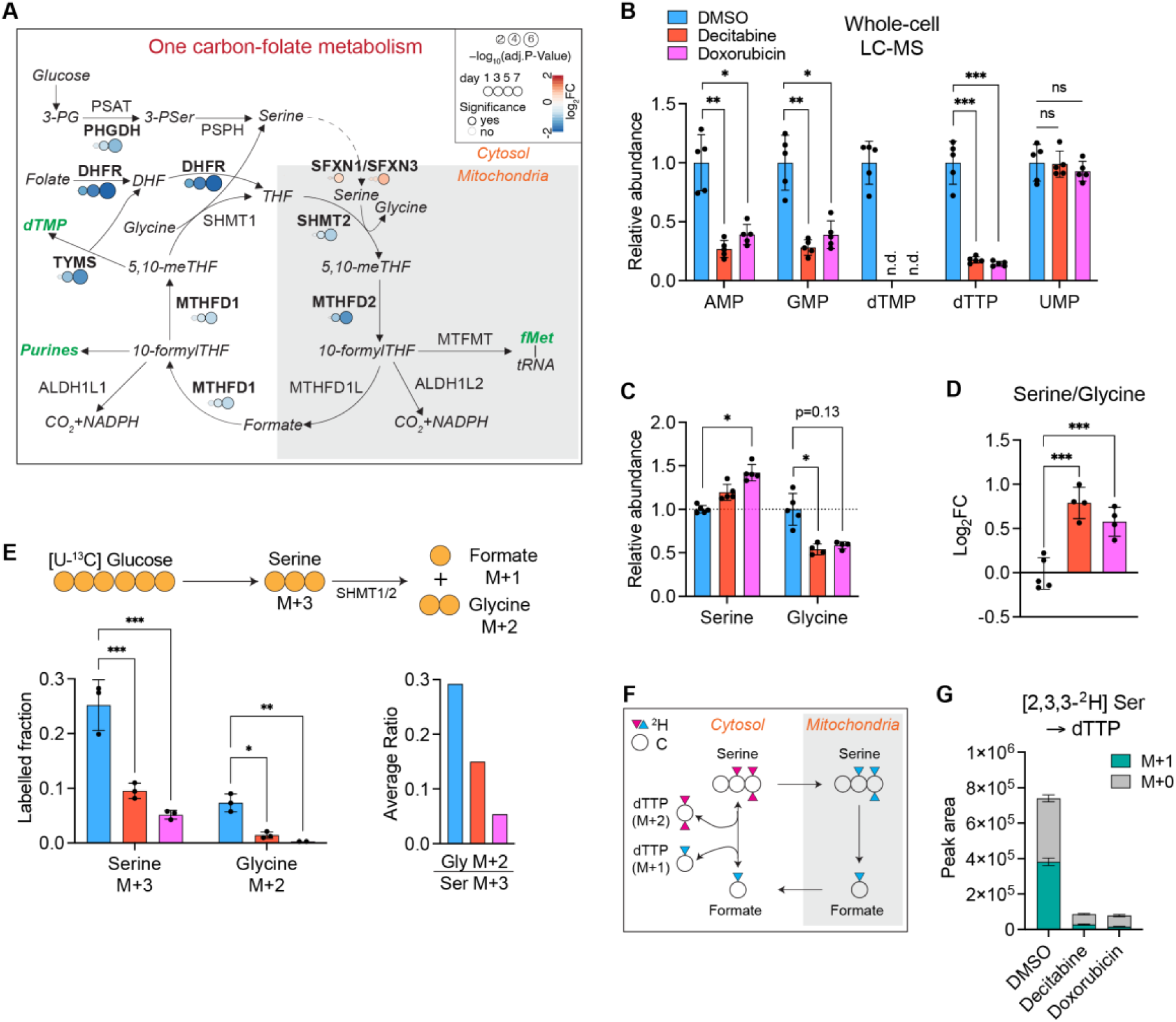
Early downregulation of 1C-folate metabolism coordinated between mitochondria and cytosol in senescent fibroblasts. (A) The mammalian 1C-folate metabolism. Enzyme levels are derived from the proteomics data. DEGs are shown in bold font. The color code indicates Log2 fold changes. The major products of the pathway are shown in green italicized font. fMet: formyl-methionine. (B) Steady-state levels of nucleotides relevant to 1C-folate metabolism are shown. The statistical analysis was performed for Log2 fold changes of each treatment compared to DMSO. See method and Supplementary figure 5. n.d.: not detected. Welch t-test, Bonferroni-Dunn correction. n=5 from independent cultures. (C, D) Steady-state levels of serine and glycine and their ratio. One outlier in the measurement of glycine was excluded from both decitabine- and doxorubicin-treated samples. One-way ANOVA, Dunnett correction. n=4-5 from independent cultures. (E) Metabolic tracing of serine and glycine using 5.5 mM [U-^13^C] glucose added to the MEM (^13^C:^12^C=1:1) for 24 h in IMR90 fibroblasts on day 7 after the treatment of DMSO, decitabine, or doxorubicin. One-way ANOVA, Dunnett correction. n=3 from independent cultures. (F) Schematic diagram of the serine metabolism highlighting the compartmentalized flux of carbons and hydrogens through the 1C-folate cycle. (G) Metabolic tracing of dTTP using [2,3,3-^2^H] serine (200 µM) for 24 h in IMR90 fibroblasts on day 7 after the treatment with DMSO, decitabine, or doxorubicin. One-way ANOVA, Dunnett correction. n=3 from independent cultures.

Accordingly, inhibition of the 1C-folate/serine catabolism in senescent cells results in the depletion of deoxythymidines in the senescent cells (Figure 6B), without any detectable increase in the cytosolic catalysis of serine (Figure 6G). Thus, the 1C-folate metabolism is downregulated in the senescent cells in accordance with the proteomic analysis.

## Discussion

We have performed a time-resolved proteomic analysis to define mitochondrial adaptations in senescent cells. Using anti-cancer drugs in human fibroblasts as a CS model (38), we show broad reshaping of the mitochondrial proteome and metabolic rewiring of mitochondria upon establishment of the senescent state. About 40% of the mitochondrial proteins were significantly changed in senescent cells, with membrane proteins being generally enriched over soluble matrix proteins. Our time-resolved proteomic analysis did not only establish a broad rewiring of the mitochondrial proteome but also allowed us to distinguish early and late proteomic adaptations. Our enrichment analysis yielded primarily metabolism-related signatures among the early responding pathways, highlighting the importance of metabolic rewiring of mitochondria in senescence.

We identified enhanced catabolism of BCAAs as an early-responding metabolic pathway in senescent cells. We observed an increased flux of both nitrogen and carbons of BCAAs to their downstream metabolites, such as some NEAAs and acyl-CoAs. On one hand, transamination of BCAAs to the NEAAs supports the maintenance of the levels of alanine, glutamate, proline, and serine, which we found to be preserved in senescent cells. Acyl-CoAs, on the other hand, are used for lipid synthesis. In agreement with previous findings showing that senescent cells enhance both synthesis and oxidation of fatty acids (13, 39–42), we observed a late upregulation of lipid metabolizing proteins in our proteomic analysis. It is therefore conceivable that BCAAs serve as a source for lipid synthesis in senescent cells, as has been described for adipose tissue (43, 44).

In contrast to BCAA degradation, the 1C-folate metabolism was downregulated at the early stages of CS development. The 1C-folate cycle provides intermediates for the synthesis of purines and dTMP in the cytosol which are required for the replication of the genome. Low demand for nucleotide synthesis with the reduced 1C-folate metabolism is therefore consistent with the stably cell-cycle arrested state of senescent cells. Indeed, inhibition of deoxynucleotide metabolism was found to be both necessary and sufficient for oncogene-induced senescence (45). In addition to its role in nucleotide synthesis, the 1C-folate cycle supports mitochondrial translation by supplying formyl-methionine for translation initiation in mitochondria (46). Our metabolic tracing experiments revealed that serine is catabolized exclusively via the mitochondrial arm of the 1C-folate cycle, which is downregulated in senescent cells without compensation from the cytosol. Accordingly, we observed reduced mitochondrial translation in senescent cells, which is consistent with the decreased bioenergetic activity of mitochondria in these cells. Since a decrease in mitochondrial translation and OXPHOS activity makes cells vulnerable to inhibition of glycolysis (46), our data would explain such susceptibility of senescent cells to glucose restriction (27). It should be noted that mitochondrial translation is a late- responding pathway upon induction of the senescent state. This likely explains why we did not observe a general decrease of OXPHOS subunits in our proteomic analysis, despite attenuated mitochondrial translation.

The mitochondrial volume was increased >12-fold on average in senescent IMR90 fibroblasts. The increased mitochondrial volume in senescent cells can be attributed to both enhanced mitochondrial biogenesis and impaired mitophagy (10). For example, the mitochondrial biogenesis factor PGC-1β mediates the increase of mitochondrial abundance in senescent cells (11). On the other hand, mitophagy was shown to be impaired in these cells (47), which also exhibit lysosomal dysfunctions (48). However, it is important to note that despite the larger mitochondrial volume, the mitochondrial proteome scaled with the cellular proteome in senescent cells which are about 8-fold larger in volume compared to proliferating cells (30). This contrasts with the nuclear proteome, whose fraction on the cellular proteome decreased, and the ER proteome, whose fraction increased in senescent cells, in agreement with a previous finding (49). The changes in mitochondrial volume upon establishment of the senescent state must be taken into account when assessing mitochondrial activities. Our results revealed that senescent cells indeed accumulate mitochondria although their bioenergetic activity is reduced. Measuring mitochondrial abundance in senescent cells, often employing two-dimensional images or ΔΨm-dependent fluorescent probes (11, 18, 19), may systematically underestimate the accumulation of mitochondria and explain conflicting observations on mitochondrial functions and fitness in these cells.

Together, our findings establish extensive reprogramming of the mitochondrial proteome and metabolism in senescent fibroblasts. We demonstrate increased BCAA degradation and lipid metabolism and decreased 1C-folate metabolism and OXPHOS activities associated with reduced mitochondrial translation in senescent cells. Since mitochondria dictate the profile of the SASP, it is conceivable that metabolic rewiring of mitochondria is required for and shapes the SASP and therefore may impact the effects of senescent cells in the context of age-related diseases such as cancer.

## Materials and Methods

### Cell culture and chemicals

Human lung IMR90 fibroblasts were obtained from ATCC (CCL-186) and maintained in Minimum Essential Medium (MEM+glutaMAX, Thermo; 41090) supplemented with 9.5% FBS (Sigma; F7524). IMR90 cells were cultured under 3% O_2_, 5% CO_2,_ and 92% N_2_. Cells with SA-β-Gal positivity less than 10% of the population were used in all experiments. Upon the induction of CS, the medium was replaced every other day to exclude nutrient availability as a limiting factor for CS. For lentivirus production, HEK293T cells were maintained in DMEM (Thermo; 61965) supplemented with 9.5% FBS. All cells were cultured without antibiotics and routinely checked for Mycoplasma contamination. The cell number was calculated with trypan blue using Countess automated cell counter (Thermo). The chemicals used in the cell culture experiments are as follows: DMSO (Sigma; D2650), decitabine (Abcam; ab120842), doxorubicin (Sigma; D1515). Decitabine and doxorubicin were dissolved in DMSO and H_2_O, respectively.

### Establishment of cellular senescence

IMR90 fibroblasts were seeded on a diverse size of culture vessels with the density of 2,100/cm^2^ for DMSO (0.01% v/v) and decitabine (1 μM), or 6,500/cm^2^ for doxorubicin (300 nM) treatment. Cells were treated with the compounds on the next day and, subsequently, the medium was replaced every other day. DMSO and decitabine were present in the media at all times, while doxorubicin was washed out after the first medium change. Unless denoted otherwise, the timing of cell harvest was synchronized to be 24 (±3) hours from the last medium replacement and DMSO-treated cells were timely re-plated so that they did not reach the confluence by the time of harvest to maintain a proliferating state. All senescence assays were performed 7 days after the initial treatment unless otherwise specified.

### Cell proliferation assay

Cells were incubated with 10 μM EdU in DMSO for 24 h corresponding to the population doubling time. Cells were collected by trypsinization and then processed according to the manufacturer’s protocol (Thermo; C10634). The number of EdU-positive cells was counted by flow cytometry (BD Biosciences; FACS Canto) in the APC channel using conventional FSC/SSC gating criteria without a viability dye.

### Measurement of mitochondrial membrane potential, superoxide, and polarized mitochondria by flow cytometry

Cells were seeded on a 6-well plate with the density described above. On day 7, cells were collected by trypsinization and pelleted, and then processed according to the manufacturer’s protocol for labeling with mitoSOX (Thermo; M36008), TMRM (Thermo; M20036), and Mitotracker Deep Red FM (Thermo; M22426). Briefly, the collected cell pellets were resuspended in the 1 ml PBS with mitoSOX (5 μM), TMRM (20 nM), or Mitotracker Deep Red FM (50 nM) and incubated in a non-CO_2_ incubator at 37°C for 20 min. Cells were pelleted and washed with PBS twice and DAPI (1 ng/ml) was added to select live cells. Then, cells were filtered through a 50 μm cell strainer and analyzed by flow cytometry (BD Biosciences; FACScanto) in the corresponding channels (PE or APC) with the conventional SSC/FSC single cell gating strategy. The mean fluorescence intensity of the gated population was taken.

### Senescence-associated β-galactosidase assay

Cells were washed twice with PBS and subject to SA-β-Gal assay according to the manufacturer’s protocol (Abcam; ab65351). On the next day, cells were washed twice with PBS and permeabilized with 0.2% TX-100/PBS for 5 min. After washing twice, DAPI (1 ng/ml) was added to allow cell counting. The images were taken under the DAPI channel and transparent channel using an EVOS microscope (Thermo). At least 50 cells per condition were analyzed.

### Quantification of mitochondrial volume

Cells were seeded and senescence was induced. DMSO-treated control cells were seeded the day before the assay was performed. On day 7, cells were washed twice with PBS and fixed with 4% PFA (Santa Cruz; sc-281692) for 15 min at room temperature (RT). After washing out PFA with PBS twice, cells were permeabilized with 0.2% TX-100 for 5 min at RT. Cells were washed twice and incubated with the antibody against ATP5B (Invitrogen; A21351; diluted 1:1000 in 1% BSA/PBS) overnight at 4°C. On the next day, the primary antibody was washed out and goat anti-mouse IgG (H+L) antibody conjugated with Alexa fluor 568 (Invitrogen; A11031) was added (1:1000 in PBS with Alexa Fluor 647 Phalloidin (Invitrogen; A22287)) for F-actin staining to identify single cells. After 1 h, DAPI (1 ng/ml) was added after washing out the secondary antibodies for 5 min and mounted on the slides (Thermo; P10144). At least one day after the mounting, the images were taken using a confocal microscope (Leica; SP8-DLS). Z-stack confocal images were taken with 0.2 μm intervals from the bottom to the top of mitochondria. After a single cell was defined in each image based on the F-actin staining using the software Fiji (50), the stacks of 2-dimensional mitochondrial images were converted into the 3-dimensional model by Mitograph 3.0 (29). The total length, average width, and volume (by length) of mitochondria per cell were calculated by MitoGraph 3.0.

### Real-time quantitative PCR (RT-qPCR)

RNA was harvested from the cells (Macherey-Nagel; 740955) and subjected to cDNA synthesis with oligo(dT) reverse transcriptase (Promega; A2791) according to the manufacturer’s protocol. Target mRNA levels were quantified by ΔΔCt values using TaqMan fast advanced master-mix (Thermo; 4444557) with the TaqMan probes as follows: B2M (Hs99999907_m1), IL1A (Hs00174092_m1), IL1B (Hs01555410_m1), IL6 (Hs00174131_m1), CDKN1A (Hs00355782_m1), CDKN2A (Hs00923894_m1), LMNB1 (Hs01059210_m1). Fold changes were calculated using B2M as a reference control.

### Quantification of mtDNA copy number difference

Cellular DNA was extracted from the cells (Qiagen; 69504) and the mtDNA copy number was measured using the TaqMan assay as described above. Genomic DNA was measured using ACTB as a probe (Hs03023880_g1) and mtDNA was measured by two different probes (MT-ND1; Hs02596873_s1 and MT-7s; Hs02596861_s1). mtDNA copy number differences were calculated (MT-ND1/ACTB or MT-7s/ACTB) and represented by MT-ND1/ACTB as both values were comparable.

### Measurement of oxygen consumption rate (OCR)

Mitochondrial respiration was measured using an XFe96 Seahorse analyzer (Agilent; 103015) according to the manufacturer’s protocol. Briefly, 2x10e4 (proliferating) and 3x10e4 (senescent) cells were seeded per well on XFe96 plate. The next day, cells were washed twice and incubated for 1 h at 37°C in the non-CO_2_ chamber with 180 μl of the assay medium (Agilent; 103575) supplemented with L-glutamine (2 mM) and D-glucose (5.5 mM). OCR was measured with subsequent injections of the following compounds (1 μM oligomycin, 0.5 μM FCCP or CCCP, and rotenone and antimycin A (0.5 μM each)). After the assay, cells were washed once with PBS and lysed in 25 μl of SDS buffer (50 mM Tris-HCl pH 7.4, 1% SDS), followed by the BCA protein quantification. OD_562nm_ value was used as the protein amount without standards. The data were first normalized to protein amounts, followed by scaling to the cell number by a cell-to-protein ratio (Supplementary figure 1G). Spare respiratory capacity and proton leak were calculated from the OCR data by the Seahorse XF report generator (Agilent).

### Measurement of the cell-to-protein ratio

On day 7 after the treatment with H_2_O, DMSO, decitabine, or doxorubicin, IMR90 fibroblasts were trypsinized and live cells were counted with trypan blue. The cells were pelleted and lysed in the SDS buffer (50 mM Tris-HCl pH 7.4, 1% SDS) and the protein mass was measured by the BCA method.

### Isolation of mitochondria

The preparation of mitochondria-enriched membrane organelle was done as described with a few modifications (51). Briefly, cells with around 80% confluence on three 15-cm dishes were collected by scraping and washed with ice-cold PBS twice. All subsequent steps were performed at 4°C. The cells were incubated for 10 min in 1 ml isolation buffer (10 mM HEPES-KOH pH 7.4, 225 mM mannitol, 75 mM sucrose, 1 mM EGTA). Then, cells were homogenized by passing 10 times through a 27G needle. The homogenates were spun down at 800x*g* for 5 min to remove cell debris. Supernatants were centrifuged at 7000x*g* for 10 min, followed by two washing steps with an isolation buffer. The final pellets containing membrane organelles without cytosolic fraction were resuspended in the 200 μl isolation buffer.

### Mitochondrial translation *in organello* assay

Equal amounts of mitochondria (100-150 μg) were resuspended in 1 ml translation buffer (60 μg/ml of each of 19 proteogenic amino acids except methionine, 5 mM ATP, 200 μM GTP, 6 mM creatine phosphate, 60 μg/ml creatine kinase, 100 mM D-mannitol, 10 mM sodium succinate dibasic hexahydrate, 80 mM KCl, 5 mM MgCl_2_ hexahydrate, 1 mM KH_2_PO_4_, 25 mM HEPES, adjusted to pH 7.4 with KOH). 17 μl of ^35^S-methionine (Hartmann Analytic; SRM-01) was added and the sample was incubated for 1 h at 37°C under gentle mixing (300 rpm). Mitochondria were pelleted at 7000x*g* for 2 min at 4°C and resuspended in translation buffer, followed by 10 min incubation at 37°C under gentle mixing (300 rpm). Mitochondria were washed 3 times with translation buffer to remove any residual ^35^S- methionine and then resuspended in 100 μl sample buffer and run on 12% Tris-tricine SDS-PAGE. The gel was transferred to a nitrocellulose membrane and dried in the air. The radioactivity was captured by the storage phosphor screen overnight and detected by the Typhoon phosphor-imager (Cytiva Lifesciences). Membranes were blocked in 5% skim milk in TBS-T (20 mM Tris-HCL pH7.4, 150 mM NaCl, 0.1% Tween 20, pH 7.4) for 1 h, washed three times with TBS-T, and incubated with primary antibodies in 5% BSA TBS-T against SDHA (1:10000, Abcam; ab14715), MT-CO1 (1:10000, Abcam; ab14705), MT-ND1 (1:2000, Abcam; ab181848), MT-CO2 (1:1000, Invitrogen; A6404), MT-ATP8 (1:2000, Proteintech; 26723-1-AP), MT-ATP6 (1:2000, Proteintech; 55313-1-AP). Then membranes were washed 3 times with TBS-T and incubated with goat anti-mouse or anti-rabbit IgG HRP-conjugated secondary antibody (1:10000 in 5% BSA TBS-T, Bio-rad; #1706515, #1706516) for 1 h, at RT. Membranes were washed again and developed using enhanced chemiluminescence and analyzed by ChemoStar Touch (INTAS Science Imaging) and the Fiji software (50).

### Proteomics: peptide preparation

Cells were seeded on 15-cm dishes and, on the next day, treated with either DMSO or decitabine. On day 1, 3, 5, and 7, cells were collected and washed twice with PBS. In all conditions, cells were collected at a confluency of around 80%. Cell pellets were resuspended in 15 µl of lysis buffer (6 M guanidinium chloride, 2.5 mM tris(2-carboxyethyl) phosphine, 10 mM chloroacetamide, 100 mM tris-hydrochloride) and heated at 95°C for 10 min. The lysates were sonicated (30 s/30 s, 10 cycles, high performance) by Bioruptor (Diagenode; B01020001), followed by centrifugation at 21000x*g* for 20 min at 20°C. 200 µg of supernatants were digested with 1 µl trypsin (Promega; V5280) overnight at 37°C. On the next day, formic acid was added to the digested peptide lysates (to 1% final concentration) to stop trypsin digestion, and samples were desalted by homemade STAGE tips (52). Eluted lysates in 60% acetonitrile/0.1% formic acid were dried by vacuum centrifugation (Eppendorf; Concentrator Plus) at 45°C.

### Proteomics: TMT labeling

4 µg of desalted peptides were labeled with tandem mass tags TMT10plex (Thermo; 90110) using a 1:20 ratio of peptides to TMT reagent. TMT labeling was carried out according to the manufacturer’s instruction with the following changes: dried peptides were reconstituted in 9 µl 0.1 M TEAB, to which 7 µl TMT reagent in acetonitrile was added to a final acetonitrile concentration of 43.75%. The reaction was quenched with 2 µl 5% hydroxylamine. Labeled peptides were pooled, dried, resuspended in 0.1% formic acid, split into two samples, and desalted using homemade STAGE tips (52).

### Proteomics: high-pH fractionation

Pooled TMT labeled peptides were separated on a 150 mm, 300 μm OD, 2 μm C18, Acclaim PepMap (Thermo) column using an Ultimate 3000 (Thermo). The column was maintained at 30°C. Separation was performed with a flow of 4 μl using a segmented gradient of buffer B from 1% to 50% for 85 min and 50% to 95% for 20 min. Buffer A was 5% acetonitrile 0.01M ammonium bicarbonate, buffer B was 80% acetonitrile 0.01 M ammonium bicarbonate. Fractions were collected every 150 s and combined into nine fractions by pooling every ninth fraction. Pooled fractions were dried in Concentrator plus (Eppendorf), and resuspended in 5 μl 0.1% formic acid, from which 2 μl were analyzed by LC-MS/MS.

### Proteomics: LC-MS/MS analysis

Dried fractions were re-suspended in 0.1% formic acid and separated on a 50 cm, 75 µm Acclaim PepMap column (Thermo; 164942) and analyzed on an Orbitrap Lumos Tribrid mass spectrometer (Thermo) equipped with a FAIMS device (Thermo). The FAIMS device was operated in two compensation voltages, -50 V and -70 V. Synchronous precursor selection based on MS3 was used for the acquisition of the TMT reporter ion signals. Peptide separation was performed on an EASY-nLC1200 using a 90 min linear gradient from 6% to 31% buffer; buffer A was 0.1% formic acid, and buffer B was 0.1% formic acid with 80% acetonitrile. The analytical column was operated at 50°C. Raw files were split based on the FAIMS compensation voltage using FreeStyle (Thermo).

### Proteomics: peptide identification and quantification

Proteomics data were analyzed using MaxQuant, version 1.5.2.8, (53). The isotope purity correction factors, provided by the manufacturer, were included in the analysis. Mitochondrial annotations were based on human MitoCarta 3.0 (34).

### Proteomics: data analysis and visualization

Differential expression analysis was performed using limma version 3.34.9 (54) and R version 3.4.3 (55). Proteins with P<0.05 (Bonferroni-Hochberg method) were deemed significant and differentially expressed. Quantified proteomics data were investigated for the enrichment analysis including statistics by the String database (56) and the GSEA (57, 58). For the GSEA analysis, the background gmt files were made with MitoPathways and localization information from the human MitoCarta 3.0. The total quantified 6482 proteins were used as a background. For the categorization of organellar proteome, the reference proteome was used from the publicly available data as described in the main text (Figure 3B). Graphs were drawn by GraphPad Prism version 9.3.1 and Supplementary figure 3 and 4 by R version 3.4.3.

### Metabolomics: metabolite preparation

Cells on 6-well plates were washed twice with the wash buffer (75 mM ammonium carbonate, pH 7.4) and the plates were flash-frozen in liquid nitrogen. 800 μl extraction buffer (acetonitrile:methanol:H_2_O=4:4:2, -20°C) was added to the wells, scraped, and centrifuged by 21,000x*g* for 20 min at 4°C. The supernatants were dried by vacuum centrifugation (Labogene) for 6 h at 20°C. Pellets were lysed in 50 mM Tris-KOH pH 8.0, 150 mM NaCl, 1% SDS, and used for protein quantification using the BCA assay (Thermo; 23225). To measure steady-state levels of metabolites, the following internal standards were added to the extraction buffer: 2.5 mM amino acids standard (CIL; MSK-A2-1.2), 100 μg/ml citrate d_4_ (Sigma; 485438), 1 mg/ml ^13^C_10_ ATP (Sigma; 710695). No internal standard was added for the isotopologue tracing experiments. Isotopologues used in the experiments are as following: ^13^C_6_ D-glucose (Sigma; 389374), 2,3,3-^2^H L-serine (CIL; DLM-582), ^13^C_6_ L-leucine (Sigma; 605239), ^15^N L-leucine (sigma; 340960), ^13^C_5_ L-valine (Sigma; 758159), ^15^N L-valine (Sigma; 490172). Isotopologues were added to the regular culture medium (MEM supplemented with 9.5% undialyzed FBS) and treated to cells as indicated in each figure legend.

### Metabolomics: Anion-Exchange Chromatography Mass Spectrometry (AEX-MS) for the analysis of anionic metabolites

Extracted metabolites were re-suspended in 200 µl of Optima UPLC/MS grade water (Thermo). After 15 min incubation on a thermomixer at 4°C and a 5 min centrifugation at 16000*xg* at 4°C, 100 µl of the cleared supernatant was transferred to polypropylene autosampler vials (Chromatography Accessories Trott). The samples were analyzed using a Dionex ion chromatography system (Integrion, Thermo) as described previously (59). In brief, 5 µl of polar metabolite extract were injected in full loop mode using an overfill factor of 1, onto a Dionex IonPac AS11-HC column (2 mm × 250 mm, 4 μm particle size, Thermo) equipped with a Dionex IonPac AG11-HC guard column (2 mm × 50 mm, 4 μm, Thermo). The column temperature was held at 30°C, while the autosampler was set to 6°C. A potassium hydroxide gradient was generated using a potassium hydroxide cartridge (Eluent Generator, Thermo), which was supplied with deionized water. The metabolite separation was carried at a flow rate of 380 µl/min, applying the following gradient conditions: 0-3 min, 10 mM KOH; 3-12 min, 10−50 mM KOH; 12-19 min, 50-100 mM KOH, 19-21 min, 100 mM KOH, 21-22 min, 100-10 mM KOH. The column was re-equilibrated at 10 mM for 8 min.

For the analysis of metabolic pool sizes, the eluting compounds were detected in negative ion mode using full scan measurements in the mass range m/z 50 – 750 on a Q-Exactive HF high-resolution MS (Thermo). The heated electrospray ionization (ESI) source settings of the mass spectrometer were: Spray voltage 3.2 kV, the capillary temperature was set to 300°C, sheath gas flow 60 AU, aux gas flow 20 AU at a temperature of 330°C and a sweep gas glow of 2 AU. The S-lens was set to a value of 60.

The semi-targeted LC-MS data analysis was performed using the TraceFinder software (Version 4.1, Thermo). The identity of each compound was validated by authentic reference compounds, which were measured at the beginning and the end of the sequence.

For data analysis, the area of the deprotonated [M-H+]- monoisotopic mass peak of each compound was extracted and integrated using a mass accuracy of <5 ppm and a retention time (RT) tolerance of <0.05 min as compared to the independently measured reference compounds. Areas of the cellular pool sizes were normalized to the internal standards added to the extraction buffer, followed by total ion counts (TIC) normalization.

### Metabolomics: semi-targeted liquid chromatography-high-resolution mass spectrometry-based (LC-HRS-MS) analysis of amine-containing metabolites

The LC-HRMS analysis of amine-containing compounds was performed using an adapted benzoylchloride-based derivatization method (60). In brief, the polar fraction of the metabolite extract was re-suspended in 200 µl of LC-MS-grade water (Optima-Grade, Thermo) and incubated at 4°C for 15 min on a thermomixer. The re-suspended extract was centrifuged for 5 min at 16000 x g at 4°C and 50 µl of the cleared supernatant was mixed with 25 µl of 100 mM sodium carbonate (Sigma), followed by the addition of 25 µl 2% [v/v] benzoylchloride (Sigma) in acetonitrile (Optima-Grade, Thermo). Samples were vortexed and kept at 20°C until analysis. For the LC-HRMS analysis, 1 µl of the derivatized sample was injected onto a 100 x 2.1 mm HSS T3 UPLC column (Waters). The flow rate was set to 400 µl/min using a binary buffer system consisting of buffer A (10 mM ammonium formate (Sigma), 0.15% [v/v] formic acid (Sigma) in LC-MS-grade water (Optima-Grade, Thermo). Buffer B consisted solely of acetonitrile (Optima-grade, Thermo). The column temperature was set to 40°C, while the LC gradient was: 0% B at 0 min, 0-15% B 0- 4.1min; 15-17% B 4.1 – 4.5 min; 17-55% B 4.5-11 min; 55-70% B 11 – 11.5 min, 70-100% B 11.5 - 13 min; B 100% 13 - 14 min; 100-0% B 14 -14.1 min; 0% B 14.1-19 min; 0% B. The mass spectrometer (Q-Exactive Plus, Thermo) was operating in positive ionization mode recording the mass range m/z 100-1000. The heated ESI source settings of the mass spectrometer were: Spray voltage 3.5 kV, capillary temperature 300°C, sheath gas flow 60 AU, aux gas flow 20 AU at a temperature of 330°C, and the sweep gas to 2 AU. The RF-lens was set to a value of 60. Semi-targeted data analysis for the samples was performed using the TraceFinder software (Version 4.1, Thermo). The identity of each compound was validated by authentic reference compounds, which were run before and after every sequence. Peak areas of [M+nBz+H]+ ions were extracted using a mass accuracy (<5 ppm) and a retention time tolerance of <0.05 min. Areas of the cellular pool sizes were normalized to the internal standards ([U]-^15^N;[U]-^13^C amino acid mix (MSK-A2-1.2), Cambridge Isotope Laboratories), which were added to the extraction buffer, followed by normalization to the TIC.

### Metabolomics: semi-targeted liquid chromatography-high-resolution mass spectrometry-based (LC-HRS-MS) analysis of Acyl-CoA metabolites

The LC-HRMS analysis of Acyl-CoAs was performed using a modified protocol based on the previous method (60). In brief, the polar fraction of the metabolite extract was re-suspended in 50 µl of LC-MS-grade water (Optima-Grade, Thermo). For the LC-HRMS analysis, 1 µl of the sample was injected onto a 30 x 2.1 mm BEH Amide UPLC column (Waters) with a 1.7 µm particle size. The flow rate was set to 500 µl/min using a quaternary buffer system consisting of buffer A (5 mM ammonium acetate, Sigma) in LC-MS-grade water (Optima-Grade, Thermo). Buffer B consisted of 5 mM ammonium acetate (Sigma) in 95% acetonitrile (Optima-grade, Thermo). Buffer C consisted of 0.1% phosphoric acid (85%, VWR) in 60% acetonitrile (acidic wash) and buffer D of 50% acetonitrile (neutral wash). The column temperature was set to 30°C, while the LC gradient was: 85% B for 1 min, 85- 70% B 1- 3min; 70-50% B 3 – 3.2 min; holding 50% B till 5 min; 100% C 5.1 – 8 min, 100% D 8.1 - 10 min; followed by re-equilibration 85% B 10.1 - 13 min. The mass spectrometer (Q-Exactive Plus, Thermo) was operating in positive ionization mode recording the mass range m/z 760-1800. The heated ESI source settings of the mass spectrometer were: Spray voltage 3.5 kV, capillary temperature 300°C, sheath gas flow 50 AU, aux gas flow 15 AU at a temperature of 350°C, and the sweep gas to 3 AU. The RF-lens was set to a value of 55.

Semi-targeted data analysis for the samples was performed using the TraceFinder software (Version 4.1, Thermo). The identity of Acetyl-CoA and Malonyl-CoA was validated by authentic ^13^C-labelled reference compounds, which were run before. Other Acyl-CoAs were validated by using *E. coli* reference material matching exact mass and reporter ions from PRM experiments. Peak areas of [M+H]+ ions and corresponding isotopomers were extracted using a mass accuracy (<5 ppm) and a retention time tolerance of <0.05 min. The Peak area was normalized by the TIC.

### Metabolomics: data analysis and visualization

The steady-state level of metabolites was normalized by the total ion counts (TIC) value. Statistical analysis of differential abundance was performed with fold changes in log2 values by the welch t-test with correction using the Bonferroni-Dunn method. For mass isotopologue experiments, the natural abundance of ^13^C was not corrected and the kinetic isotope effect of the ^2^H tracer was not considered. All the statistical analysis and graphs were done by GraphPad Prism version 9.3.1. For the heatmap in Supplementary figure 5C, Flaski was used (61).

### Data analysis and statistics

All statistical analyses were performed by GraphPad Prism version 9.3.1 except proteomics data. When two groups were compared, the Welch t-test was used with a multiple comparison correction by the Bonferroni-Dunn method, if needed. When 3 or more groups were compared, the ordinary ANOVA test was used. One-way ANOVA was used for multiple groups under one condition and two-way ANOVA for multiple groups under two conditions. Each subject group was compared to the control group with a multiple comparison correction by the Dunnett method. *:p<0.05, **:p<0.01, ***:p<0.001.

---

**Supplementary figure 1.**
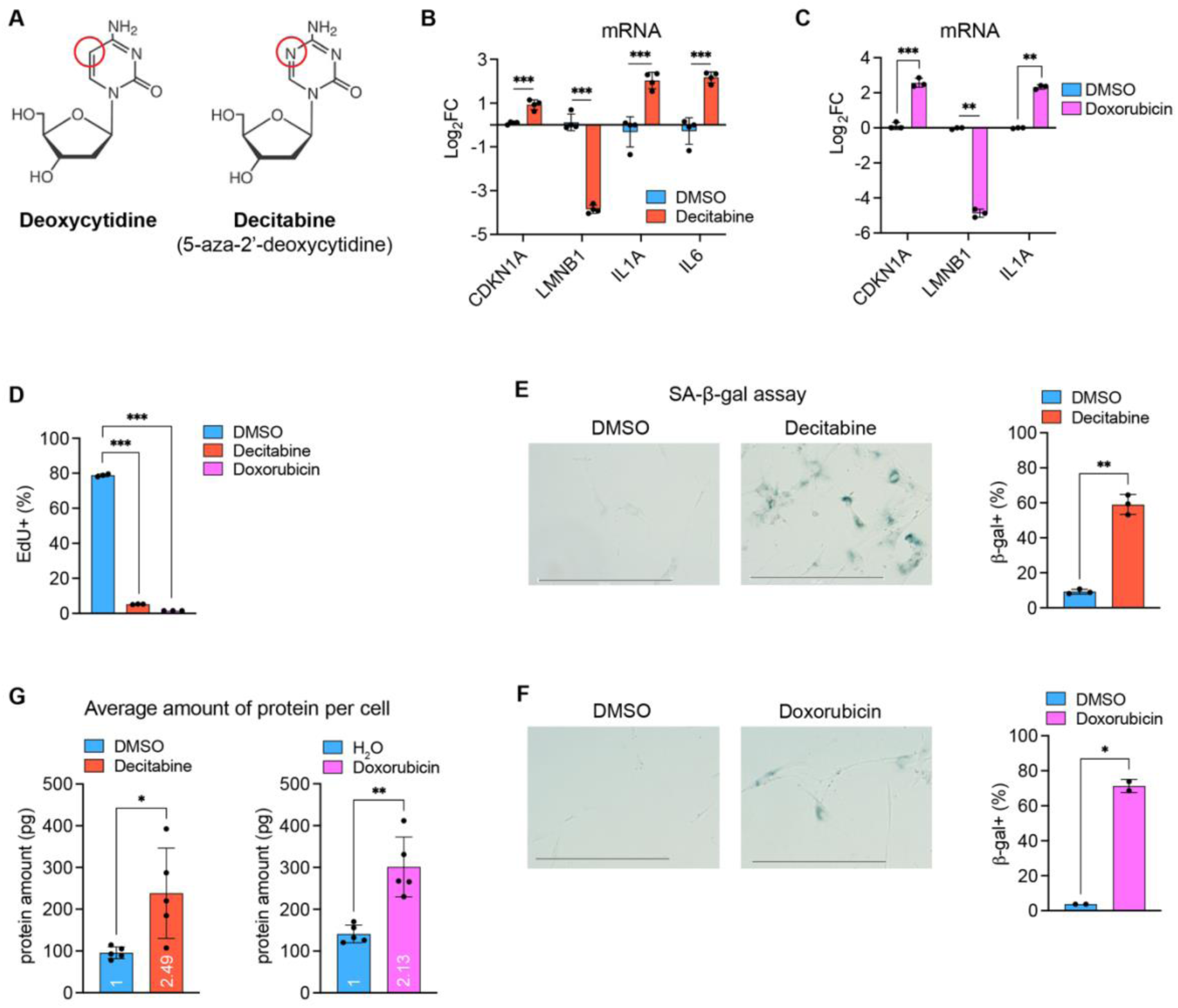
Establishment of CS in IMR90 fibroblasts. (A) Chemical structures of deoxycytidine and decitabine. (B-G) IMR90 cells were treated with DMSO (0.01%), decitabine (1 µM), or doxorubicin (300 nM) for 7 days and analyzed. (B, C) mRNA levels were measured on day 7 by RT-qPCR. Welch t-test, Bonferroni-Dunn correction. (B) n=4, (C) n=3 from independent cultures. (D) Cells were treated with EdU (10 µM) for 24 h on day 7. EdU positivity was measured by flow cytometry. One-way ANOVA, Dunnett correction. n=3 from independent cultures. (E, F) Senescence-associated β-galactosidase (SA-β-gal) assay on day 7. At least 50 cells were counted per replicate in each condition. Scale bar: 500 µm. Welch t-test. (E) n=3, (F) n=2 from independent cultures. (G) Protein mass per cell was measured on day 7. The mean fold changes are shown within the bars. Welch t-test, n=5 from independent cultures.

**Supplementary figure 2.**
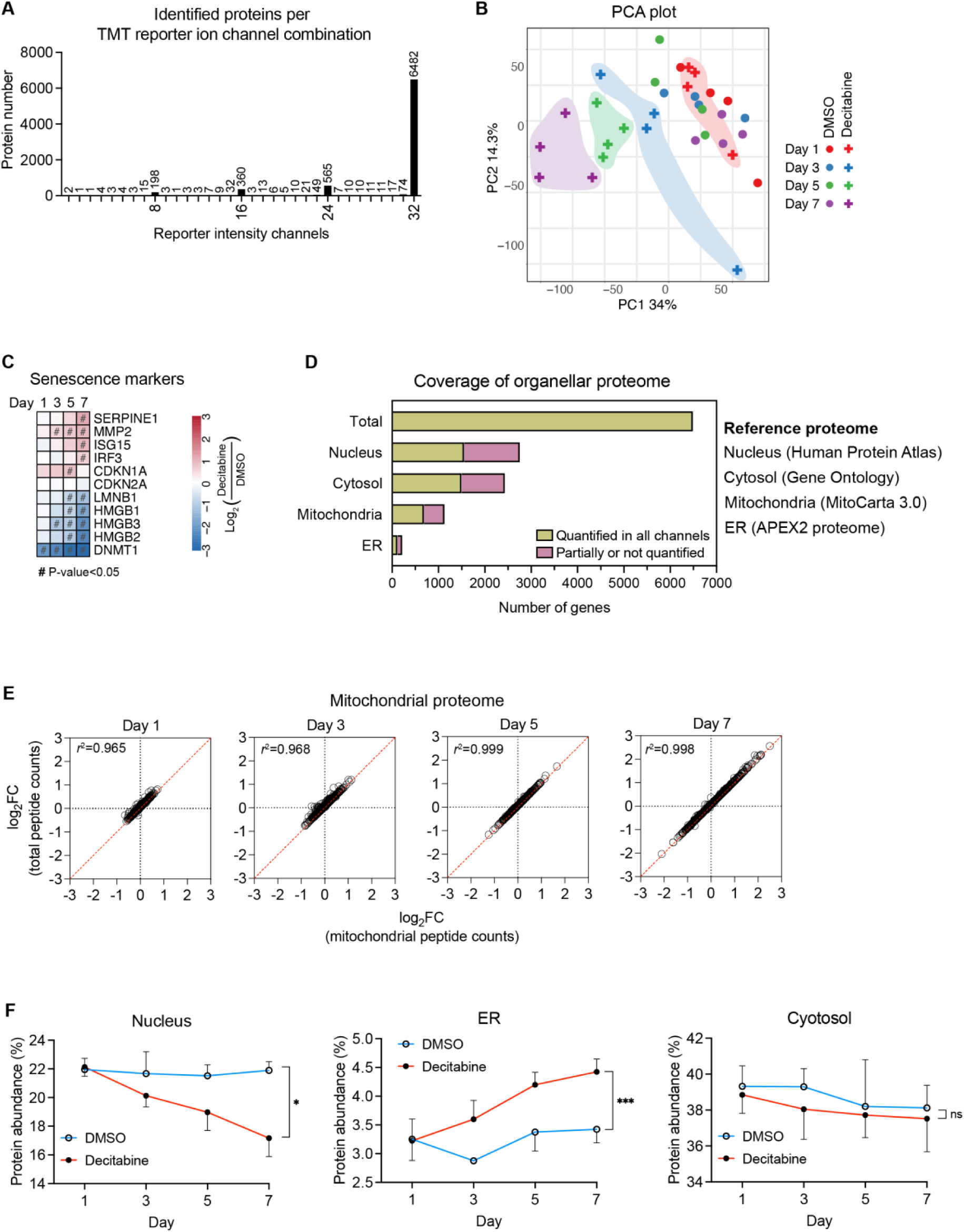
Time-resolved proteomics throughout the development of CS. (A) The number of proteins identified in 32 TMT reporter ion channels is shown. In addition, proteins only identified in groups of 8, 16, 24, and 32 TMT reporter ion channels are shown. Proteins identified in all 32 channels are subjected to subsequent analyses. (B) Principal component analysis (PCA) of the time-resolved proteome dataset. Decitabine- treated samples on each day are grouped with colors. (C) Steady-state levels of several senescence marker proteins upon the CS induction. (D) Coverage of several organellar proteomes. Reference proteomes were used as backgrounds as described in the main text for Figure 3B. Proteins quantified in all 32 samples were compared to the reference proteomes to calculate the coverage. (E) Correlation between mitochondrial proteomic change quantified by two different normalization units: total peptide counts and mitochondrial peptide counts. *r*=Pearson coefficient. (F) The protein abundance of indicated organelle was calculated as described in Figure 3B, based on the reference proteome from (D).

**Supplementary figure 3.**
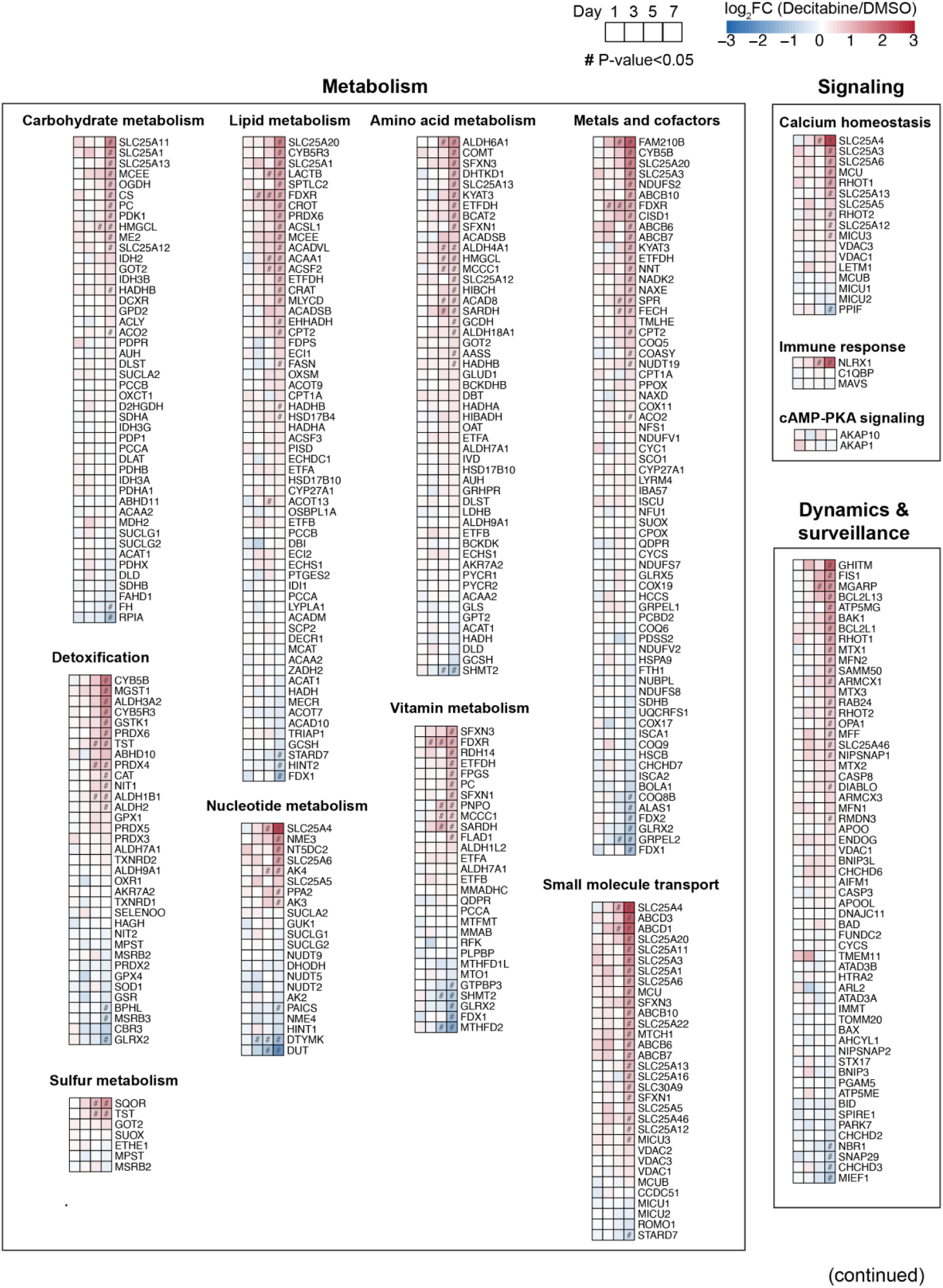
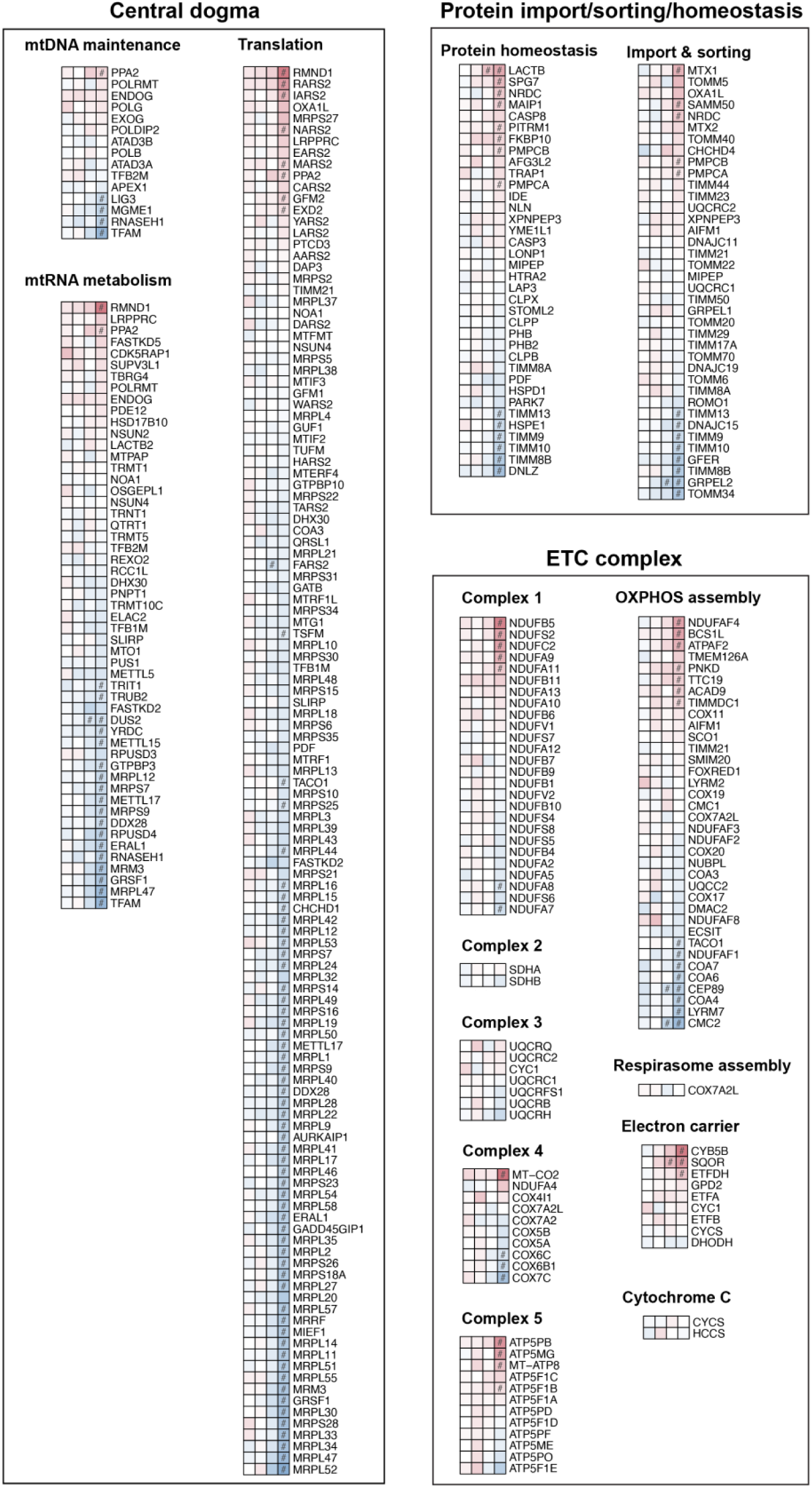
List of mitochondrial proteins quantified. All mitochondrial proteins quantified in the proteomic dataset are presented according to the categories based on the MitoCarta 3.0. Genes in each category are shown in an ascending order based on the Log_2_FC values on day 7.

**Supplementary figure 4.**
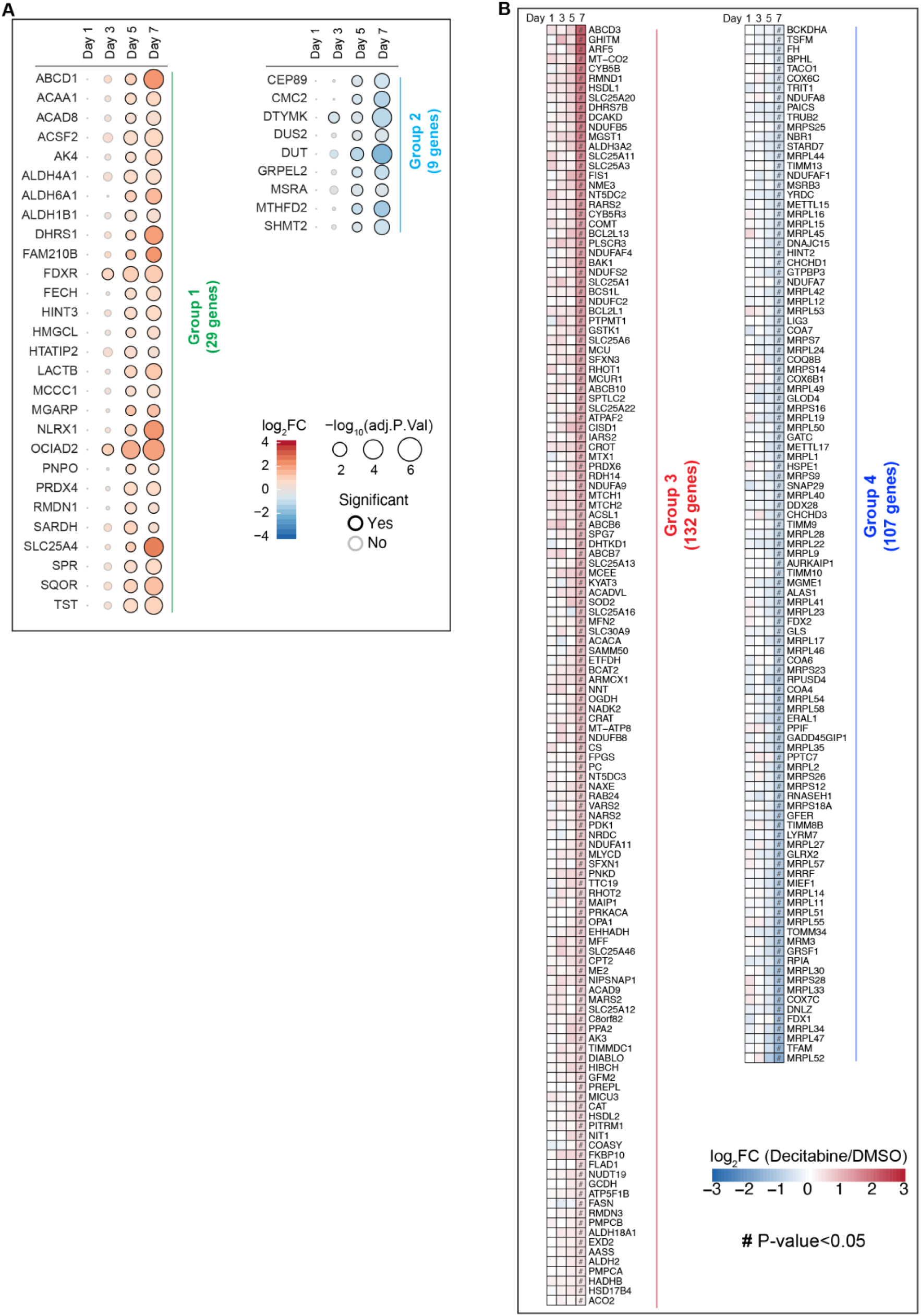
Classification of proteins according to the temporal dynamics of their steady-state levels upon CS induction. Mitochondrial proteins in early responding pathways (A, groups 1 and 2) and late responding pathways (B, groups 3 and 4) are shown.

**Supplementary figure 5.**
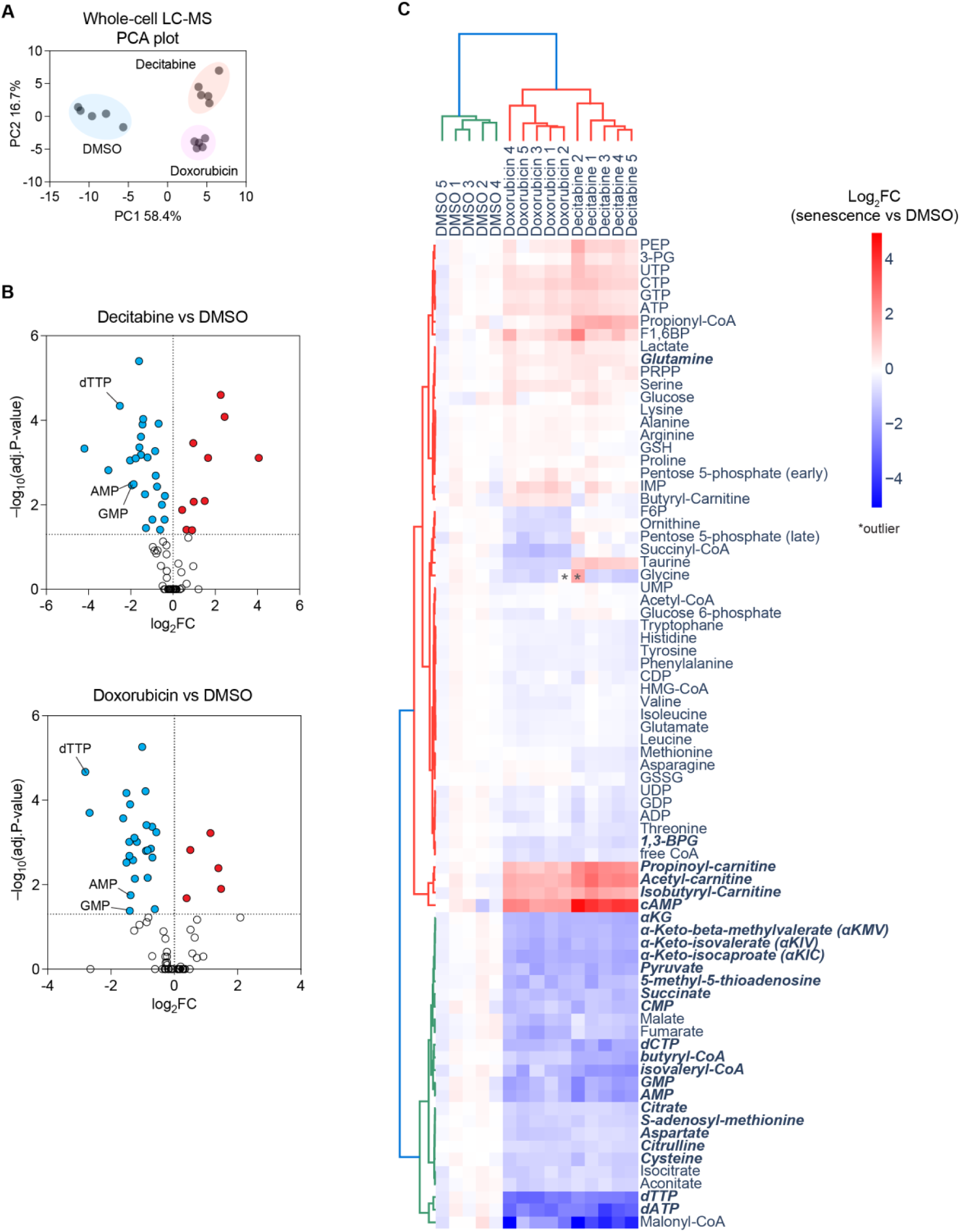
Metabolomic analysis of senescent fibroblasts on day 7 after treatment with decitabine and doxorubicin. (A) PCA plot of targeted metabolomics data measured by LC-MS. 79 metabolites were quantified in total. The peak area of each metabolite was normalized by total ion counts (TIC) and subjected to PCA. n=5 from independent cultures. (B) Volcano plots of metabolomics data. The TIC normalized peak area of each metabolite was compared between senescence and proliferating (DMSO) conditions. Significantly changed metabolites (P<0.05) are highlighted with colors. Metabolites derived from the 1C-folate metabolism are denoted. Welch t-test, Bonferroni-Dunn correction. n=5 (C) Heatmap of metabolomics data with Log_2_FC values. Treatments and metabolites are hierarchically clustered according to the Ward method with Euclidean distance. Significantly changed metabolites (P<0.05) in the same direction in both decitabine and doxorubicin senescent cells are shown in bold italicized font. Metabolites that were not detected in all samples (i.e. dTMP) are not shown.

## References

1. Childs BG, Durik M, Baker DJ, van Deursen JM. Cellular senescence in aging and age-related disease: from mechanisms to therapy. Nat Med. 2015;21(12):1424–35.

2. Basisty N, Kale A, Jeon OH, Kuehnemann C, Payne T, Rao C, et al. A proteomic atlas of senescence-associated secretomes for aging biomarker development. PLoS Biol. 2020;18(1):e3000599.

3. Demaria M, Ohtani N, Youssef SA, Rodier F, Toussaint W, Mitchell JR, et al. An essential role for senescent cells in optimal wound healing through secretion of PDGF-AA. Dev Cell. 2014;31(6):722–33.

4. Jun JI, Lau LF. The matricellular protein CCN1 induces fibroblast senescence and restricts fibrosis in cutaneous wound healing. Nat Cell Biol. 2010;12(7):676–85.

5. Krizhanovsky V, Yon M, Dickins RA, Hearn S, Simon J, Miething C, et al. Senescence of activated stellate cells limits liver fibrosis. Cell. 2008;134(4):657–67.

6. Munoz-Espin D, Canamero M, Maraver A, Gomez-Lopez G, Contreras J, Murillo-Cuesta S, et al. Programmed cell senescence during mammalian embryonic development. Cell. 2013;155(5):1104–18.

7. Storer M, Mas A, Robert-Moreno A, Pecoraro M, Ortells MC, Di Giacomo V, et al. Senescence is a developmental mechanism that contributes to embryonic growth and patterning. Cell. 2013;155(5):1119–30.

8. Baker DJ, Childs BG, Durik M, Wijers ME, Sieben CJ, Zhong J, et al. Naturally occurring p16(Ink4a)-positive cells shorten healthy lifespan. Nature. 2016;530(7589):184–9.

9. Zhang L, Pitcher LE, Yousefzadeh MJ, Niedernhofer LJ, Robbins PD, Zhu Y. Cellular senescence: a key therapeutic target in aging and diseases. J Clin Invest. 2022;132(15).

10. Dalle Pezze P, Nelson G, Otten EG, Korolchuk VI, Kirkwood TB, von Zglinicki T, et al. Dynamic modelling of pathways to cellular senescence reveals strategies for targeted interventions. PLoS Comput Biol. 2014;10(8):e1003728.

11. Correia-Melo C, Marques FD, Anderson R, Hewitt G, Hewitt R, Cole J, et al. Mitochondria are required for pro-ageing features of the senescent phenotype. EMBO J. 2016;35(7):724–42.

12. Vizioli MG, Liu T, Miller KN, Robertson NA, Gilroy K, Lagnado AB, et al. Mitochondria-to-nucleus retrograde signaling drives formation of cytoplasmic chromatin and inflammation in senescence. Genes Dev. 2020;34(5-6):428–45.

13. Quijano C, Cao L, Fergusson MM, Romero H, Liu J, Gutkind S, et al. Oncogene- induced senescence results in marked metabolic and bioenergetic alterations. Cell Cycle. 2012;11(7):1383–92.

14. Kaplon J, Zheng L, Meissl K, Chaneton B, Selivanov VA, Mackay G, et al. A key role for mitochondrial gatekeeper pyruvate dehydrogenase in oncogene-induced senescence. Nature. 2013;498(7452):109–12.

15. Nacarelli T, Lau L, Fukumoto T, Zundell J, Fatkhutdinov N, Wu S, et al. NAD(+) metabolism governs the proinflammatory senescence-associated secretome. Nat Cell Biol. 2019;21(3):397–407.

16. Wiley CD, Velarde MC, Lecot P, Liu S, Sarnoski EA, Freund A, et al. Mitochondrial Dysfunction Induces Senescence with a Distinct Secretory Phenotype. Cell Metab. 2016;23(2):303–14.

17. Miwa S, Kashyap S, Chini E, von Zglinicki T. Mitochondrial dysfunction in cell senescence and aging. J Clin Invest. 2022;132(13).

18. Passos JF, Nelson G, Wang C, Richter T, Simillion C, Proctor CJ, et al. Feedback between p21 and reactive oxygen production is necessary for cell senescence. Mol Syst Biol. 2010;6:347.

19. Passos JF, Saretzki G, Ahmed S, Nelson G, Richter T, Peters H, et al. Mitochondrial dysfunction accounts for the stochastic heterogeneity in telomere-dependent senescence. PLoS Biol. 2007;5(5):e110.

20. Park YY, Lee S, Karbowski M, Neutzner A, Youle RJ, Cho H. Loss of MARCH5 mitochondrial E3 ubiquitin ligase induces cellular senescence through dynamin- related protein 1 and mitofusin 1. J Cell Sci. 2010;123(Pt 4):619–26.

21. Lee S, Jeong SY, Lim WC, Kim S, Park YY, Sun X, et al. Mitochondrial fission and fusion mediators, hFis1 and OPA1, modulate cellular senescence. J Biol Chem. 2007;282(31):22977–83.

22. Hutter E, Renner K, Pfister G, Stockl P, Jansen-Durr P, Gnaiger E. Senescence- associated changes in respiration and oxidative phosphorylation in primary human fibroblasts. Biochem J. 2004;380(Pt 3):919–28.

23. Moiseeva O, Bourdeau V, Roux A, Deschenes-Simard X, Ferbeyre G. Mitochondrial dysfunction contributes to oncogene-induced senescence. Mol Cell Biol. 2009;29(16):4495–507.

24. Hubackova S, Davidova E, Rohlenova K, Stursa J, Werner L, Andera L, et al. Selective elimination of senescent cells by mitochondrial targeting is regulated by ANT2. Cell Death Differ. 2019;26(2):276–90.

25. Johmura Y, Yamanaka T, Omori S, Wang TW, Sugiura Y, Matsumoto M, et al. Senolysis by glutaminolysis inhibition ameliorates various age-associated disorders. Science. 2021;371(6526):265–70.

26. Dou X, Long Q, Liu S, Zou Y, Fu D, Chen X, et al. Senescent cells develop PDK4- dependent hypercatabolism and form an acidic microenvironment to drive cancer resistance. bioRxiv. 2022:2022.08.29.505761.

27. Dorr JR, Yu Y, Milanovic M, Beuster G, Zasada C, Dabritz JH, et al. Synthetic lethal metabolic targeting of cellular senescence in cancer therapy. Nature. 2013;501(7467):421-5.

28. Keij JF, Bell-Prince C, Steinkamp JA. Staining of mitochondrial membranes with 10- nonyl acridine orange, MitoFluor Green, and MitoTracker Green is affected by mitochondrial membrane potential altering drugs. Cytometry. 2000;39(3):203–10.

29. Viana MP, Lim S, Rafelski SM. Quantifying mitochondrial content in living cells. Methods Cell Biol. 2015;125:77–93.

30. Neurohr GE, Terry RL, Lengefeld J, Bonney M, Brittingham GP, Moretto F, et al. Excessive Cell Growth Causes Cytoplasm Dilution And Contributes to Senescence. Cell. 2019;176(5):1083–97 e18.

31. Thul PJ, Akesson L, Wiking M, Mahdessian D, Geladaki A, Ait Blal H, et al. A subcellular map of the human proteome. Science. 2017;356(6340).

32. Ashburner M, Ball CA, Blake JA, Botstein D, Butler H, Cherry JM, et al. Gene ontology: tool for the unification of biology. The Gene Ontology Consortium. Nat Genet. 2000;25(1):25–9.

33. Hung V, Lam SS, Udeshi ND, Svinkina T, Guzman G, Mootha VK, et al. Proteomic mapping of cytosol-facing outer mitochondrial and ER membranes in living human cells by proximity biotinylation. Elife. 2017;6.

34. Rath S, Sharma R, Gupta R, Ast T, Chan C, Durham TJ, et al. MitoCarta3.0: an updated mitochondrial proteome now with sub-organelle localization and pathway annotations. Nucleic Acids Res. 2021;49(D1):D1541–D7.

35. Wiel C, Lallet-Daher H, Gitenay D, Gras B, Le Calve B, Augert A, et al. Endoplasmic reticulum calcium release through ITPR2 channels leads to mitochondrial calcium accumulation and senescence. Nat Commun. 2014;5:3792.

36. McCann MR, George De la Rosa MV, Rosania GR, Stringer KA. L-Carnitine and Acylcarnitines: Mitochondrial Biomarkers for Precision Medicine. Metabolites. 2021;11(1).

37. Ducker GS, Chen L, Morscher RJ, Ghergurovich JM, Esposito M, Teng X, et al. Reversal of Cytosolic One-Carbon Flux Compensates for Loss of the Mitochondrial Folate Pathway. Cell Metab. 2016;23(6):1140–53.

38. Ewald JA, Desotelle JA, Wilding G, Jarrard DF. Therapy-induced senescence in cancer. J Natl Cancer Inst. 2010;102(20):1536–46.

39. Wiley CD, Sharma R, Davis SS, Lopez-Dominguez JA, Mitchell KP, Wiley S, et al. Oxylipin biosynthesis reinforces cellular senescence and allows detection of senolysis. Cell Metab. 2021;33(6):1124–36 e5.

40. Flor AC, Wolfgeher D, Wu D, Kron SJ. A signature of enhanced lipid metabolism, lipid peroxidation and aldehyde stress in therapy-induced senescence. Cell Death Discov. 2017;3:17075.

41. Millner A, Lizardo DY, Atilla-Gokcumen GE. Untargeted Lipidomics Highlight the Depletion of Deoxyceramides during Therapy-Induced Senescence. Proteomics. 2020;20(10):e2000013.

42. Fafian-Labora J, Carpintero-Fernandez P, Jordan SJD, Shikh-Bahaei T, Abdullah SM, Mahenthiran M, et al. FASN activity is important for the initial stages of the induction of senescence. Cell Death Dis. 2019;10(4):318.

43. Green CR, Wallace M, Divakaruni AS, Phillips SA, Murphy AN, Ciaraldi TP, et al. Branched-chain amino acid catabolism fuels adipocyte differentiation and lipogenesis. Nat Chem Biol. 2016;12(1):15–21.

44. Wallace M, Green CR, Roberts LS, Lee YM, McCarville JL, Sanchez-Gurmaches J, et al. Enzyme promiscuity drives branched-chain fatty acid synthesis in adipose tissues. Nat Chem Biol. 2018;14(11):1021–31.

45. Aird KM, Zhang G, Li H, Tu Z, Bitler BG, Garipov A, et al. Suppression of nucleotide metabolism underlies the establishment and maintenance of oncogene-induced senescence. Cell Rep. 2013;3(4):1252–65.

46. Minton DR, Nam M, McLaughlin DJ, Shin J, Bayraktar EC, Alvarez SW, et al. Serine Catabolism by SHMT2 Is Required for Proper Mitochondrial Translation Initiation and Maintenance of Formylmethionyl-tRNAs. Mol Cell. 2018;69(4):610–21 e5.

47. Ahmad T, Sundar IK, Lerner CA, Gerloff J, Tormos AM, Yao H, et al. Impaired mitophagy leads to cigarette smoke stress-induced cellular senescence: implications for chronic obstructive pulmonary disease. FASEB J. 2015;29(7):2912–29.

48. Tai H, Wang Z, Gong H, Han X, Zhou J, Wang X, et al. Autophagy impairment with lysosomal and mitochondrial dysfunction is an important characteristic of oxidative stress-induced senescence. Autophagy. 2017;13(1):99–113.

49. Lanz MC, Zatulovskiy E, Swaffer MP, Zhang L, Ilerten I, Zhang S, et al. Increasing cell size remodels the proteome and promotes senescence. Mol Cell. 2022;82(17):3255–69 e8.

50. Schindelin J, Arganda-Carreras I, Frise E, Kaynig V, Longair M, Pietzsch T, et al. Fiji: an open-source platform for biological-image analysis. Nat Methods. 2012;9(7):676–82.

51. Frezza C, Cipolat S, Scorrano L. Organelle isolation: functional mitochondria from mouse liver, muscle and cultured fibroblasts. Nat Protoc. 2007;2(2):287–95.

52. Rappsilber J, Ishihama Y, Mann M. Stop and go extraction tips for matrix-assisted laser desorption/ionization, nanoelectrospray, and LC/MS sample pretreatment in proteomics. Anal Chem. 2003;75(3):663–70.

53. Cox J, Mann M. MaxQuant enables high peptide identification rates, individualized p.p.b.-range mass accuracies and proteome-wide protein quantification. Nat Biotechnol. 2008;26(12):1367–72.

54. Ritchie ME, Phipson B, Wu D, Hu Y, Law CW, Shi W, et al. limma powers differential expression analyses for RNA-sequencing and microarray studies. Nucleic Acids Res. 2015;43(7):e47.

55. R Core Team. R: A Language and Environment for Statistical Computing. R Foundation for Statistical Computing; 2017.

56. Szklarczyk D, Gable AL, Nastou KC, Lyon D, Kirsch R, Pyysalo S, et al. The STRING database in 2021: customizable protein-protein networks, and functional characterization of user-uploaded gene/measurement sets. Nucleic Acids Res. 2021;49(D1):D605–D12.

57. Subramanian A, Tamayo P, Mootha VK, Mukherjee S, Ebert BL, Gillette MA, et al. Gene set enrichment analysis: a knowledge-based approach for interpreting genome- wide expression profiles. Proc Natl Acad Sci U S A. 2005;102(43):15545–50.

58. Liao Y, Wang J, Jaehnig EJ, Shi Z, Zhang B. WebGestalt 2019: gene set analysis toolkit with revamped UIs and APIs. Nucleic Acids Res. 2019;47(W1):W199–W205.

59. Schwaiger M, Rampler E, Hermann G, Miklos W, Berger W, Koellensperger G. Anion-Exchange Chromatography Coupled to High-Resolution Mass Spectrometry: A Powerful Tool for Merging Targeted and Non-targeted Metabolomics. Anal Chem. 2017;89(14):7667–74.

60. Wong JM, Malec PA, Mabrouk OS, Ro J, Dus M, Kennedy RT. Benzoyl chloride derivatization with liquid chromatography-mass spectrometry for targeted metabolomics of neurochemicals in biological samples. J Chromatogr A. 2016;1446:78–90.

61. Iqbal A, Duitama, C., Metge, F., Rosskopp, D., Boucas, J. Flaski. 2021.

